# The concentration of myelin debris-like particles in the normal rat optic nerve

**DOI:** 10.1101/2025.03.19.643597

**Authors:** June Kawano

## Abstract

In the present study, we report that the concentration of myelin debris-like myelin basic protein-immunoreactive particles was observed in the distal (anterior)-most part of the myelinated region in the normal rat optic nerve. These particles were visualized using fluorescent immunohistochemistry with a mouse monoclonal anti-human myelin basic protein (MBPh) antibody (clone SMI-99). Fluorescent double immunohistochemistry, employing both the rat monoclonal anti-cow myelin basic protein (MBPc) antibody (clone 12) and the anti-MBPh antibody, revealed that the myelin basic protein immunoreactive-particles detected by the anti-MBPc antibody nearly completely overlapped with those immunostained by the anti-MBPh antibody. Since these antibodies target different sites, it can be concluded that these particles contain authentic myelin basic protein. We hypothesized that the MBPh-immunoreactive particles represent myelin debris-like structures in the normal rat optic nerve. Quantitative morphological analyses indicated that only 2 out of 6 differences in size and shape descriptors between the particles and the myelin debris observed in the damaged optic nerve of the glaucoma rat were statistically significant. Glial fibrillary acidic protein-immunoreactivity and glutamine synthetase-immunoreactivity were observed in the particles. Most of these particles were isolated from ionized calcium-binding adapter molecule 1-labeled microglia. These findings demonstrate that the myelin debris-like MBPh-immunoreactive particles are concentrated in the distal-most part of the myelinated region. This evidence suggests that the distal-most part is under physiologically stressed conditions. Furthermore, these findings may provide valuable insights into the pathophysiological mechanisms that induce localized vulnerability of the myelin sheaths.

**Key points:**

- This article demonstrates that myelin basic protein-immunoreactive particles are densely distributed in the distal (anterior)-most part of the myelinated region in the normal rat optic nerve.
- These particles exhibit morphological characteristics akin to myelin debris observed in the damaged optic nerve of the glaucoma rat. (45 words)

**Graphical Abstract Image:** 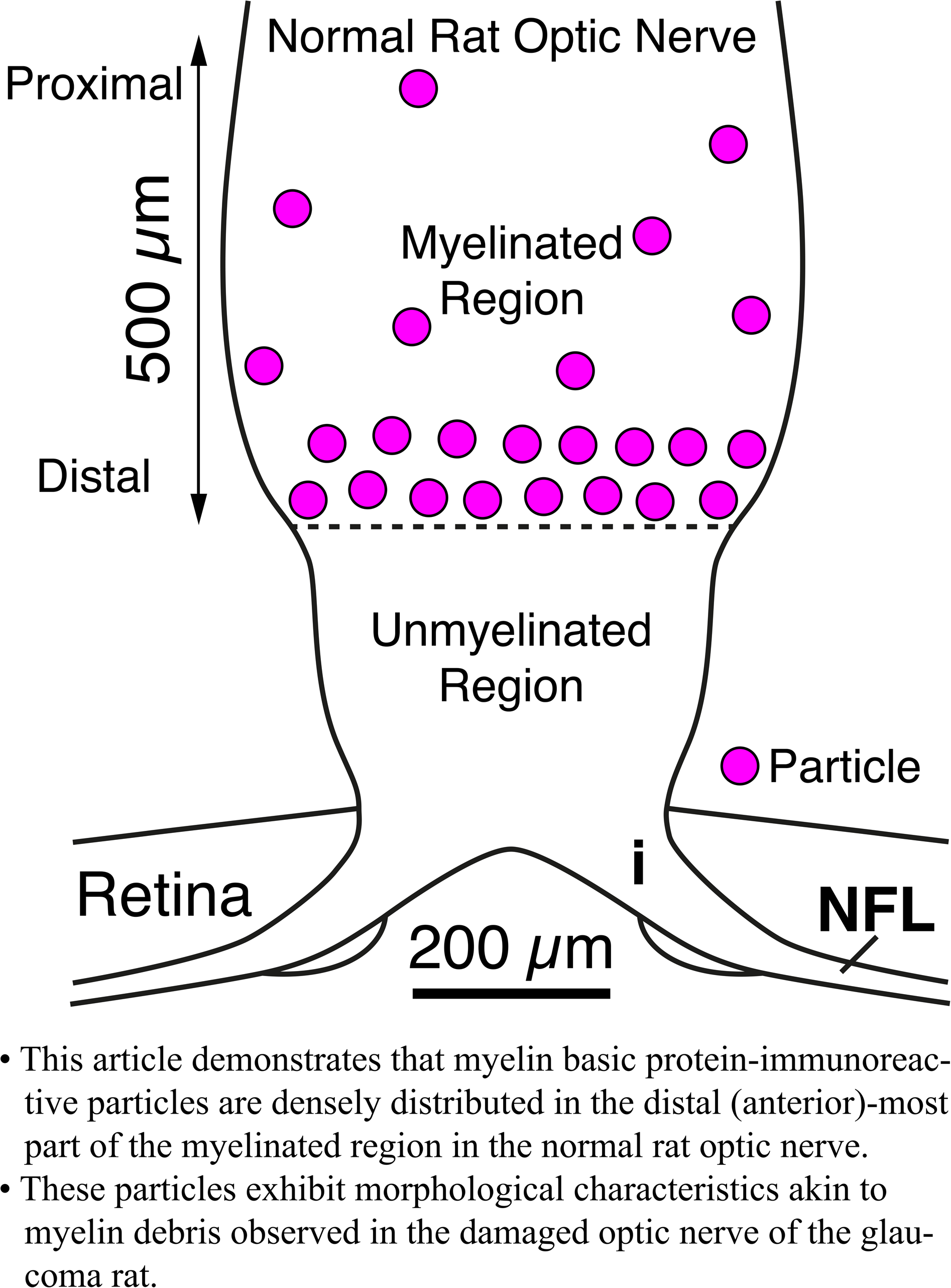

## 1 Introduction

In the myelinated region of the normal rat optic nerve, neuronal fibers are encircled by myelin sheaths except at the nodes of Ranvier (Black et al. 1985; Peters et al. 1991; Kawano 2015b). The modal diameter of the myelinated axon is 0.9 µm, while the modal diameter excluding the myelin sheath is 0.7 µm (Forrester and Peters 1967). The mean myelin thickness of large axons is 0.12 µm, whereas that of small axons is 0.08 µm (Melo et al. 2006). Individual oligodendrocytes provide, on average, 16 near axons with single myelin segments, each about 200 µm in length (Butt and Ransom 1993). Within the domain of myelin biochemistry, the principal constituents of myelin are lipids (70% of its dry weight) and proteins (30% of the dry weight). The major myelin proteins in the central nervous system are proteolipid protein (PLP, 50% of myelin protein) and myelin basic protein (MBP, 30% of myelin protein; Chrast et al. 2011; Butt 2013; Duncan and Radcliff 2016). Until recently, a traditional image (Figure 1B), characterized by the background structures and/or substances described previously, have been observed in the distal (anterior)-most part of the myelinated region. Notably, a new image (Figure 1A) demonstrating a concentration of small particles were obtained in this part. These particles were visualized using fluorescent immunohistochemistry with a mouse monoclonal anti-human myelin basic protein (MBPh) antibody (clone SMI-99; Covance, Princeton, NJ; Cat# SMI-99P; RRID: AB_10120129) as the primary antibody. Since these particles morphologically resembled the myelin debris distributed at the neural injury site, we hypothesized that the MBPh-immunoreactive particles were myelin debris-like structures in the normal rat optic nerve.

**FIGURE 1.**
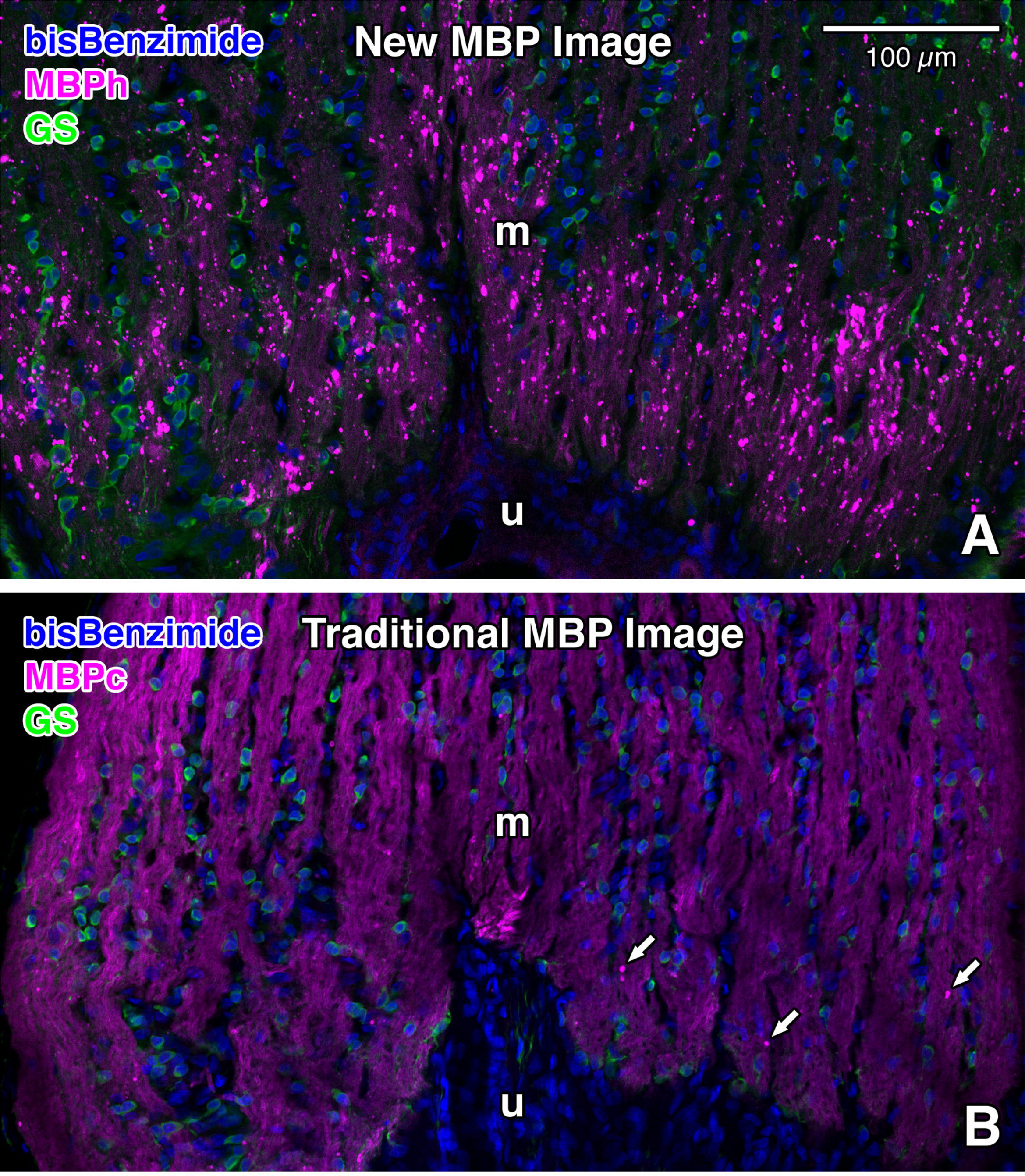
New (A) and traditional (B) images of myelin basic protein (MBP) visualized by using fluorescent double immunohistochemistry in the myelinated region of the normal rat optic nerve. Both images show sections of longitudinal slices through the paramedian part. **A:** A considerable number of particles strongly labeled in magenta are distributed in the distal (anterior)-most part of the myelinated region. These particles and myelinated nerve fibers were labeled with a mouse monoclonal anti-human myelin basic protein (MBPh) antibody (clone SMI-99; Covance, Princeton, NJ, USA; Alexa Fluor 594 label; magenta). This antibody reacts with Ala-Ser-Asp-Tyr-Lys-Ser (ASDYKS) in position 131-136 of the classic human myelin basic protein. **B:** An adjacent optic nerve section of panel **A** was double-immunostained for myelin basic protein. Myelinated nerve fibers were labeled with a rat monoclonal anti-cow myelin basic protein (MBPc) antibody (clone 12; Abcam, Cambridge, United Kingdom; Alexa Fluor 594 label; magenta). This antibody reacts with Asp-Glu-Asn-Pro-Val-Val (DENPVV) in position 82-87 of the full-length cow myelin basic protein. The arrows indicate particles strongly labeled in magenta. **A-B:** Glial cells, the majority of which were oligodendrocytes (Kawano 2015b), were immunostained with anti-glutamine synthetase (GS) antibody (Sigma-Aldrich, Saint Louis, MO, USA; Alexa Fluor 488 label; green). Cell nuclei were labeled with bisBenzimide (Hoechst 33258; blue). These two images were taken with an LSM 700 confocal microscope (Carl Zeiss, Jena, Germany). Note that the particles are concentrated in the distal-most part of the myelinated region in the normal rat optic nerve. Interestingly, a large number of the particles are seen in panel **A**; however, a smaller number are detected in a similar part of the myelinated region in panel **B**. Additionally, the majority of particles are distributed on MBP-immunoreactive myelinated nerve fibers (**A**, **B**). m, myelinated region; u, unmyelinated region. Scale bar = 100 µm in the upper right corner of **A** for **B**.

Myelin debris is produced by the breakdown of the myelin sheath immediately after neural injury. This debris persists at the injury site and contributes to regeneration failure since it contains molecules that potently inhibit axon regeneration (Chen et al. 2000; Filbin 2003) and remyelination (Kotter et al. 2006; Syed et al. 2016). Moreover, the myelin debris mediates a persistent inflammatory response during the progression of the injury (Jeon et al. 2008; Sun et al. 2010; Wang et al. 2015; Zhou et al. 2019). Thus, these findings suggest that the distal-most part, where the myelin debris-like MBPh-immunoreactive particles were concentrated, is not a physiological site, but a specific site in the myelinated region of the normal rat optic nerve. In this specific site, concentrations of GFAP (glial fibrillary acidic protein) and GS (glutamine synthetase) have been documented (Kawano 2026). The concentration of GFAP suggests reactive astrogliosis, which is a secondary event to insults in the central nervous system (Messam et al. 2002; McAteer and Choudhury 2009; Sofroniew 2009). A significant increase in GS immunoreactivity of GS-immunoreactive cells indicates a chronic pathological condition (Ben Haim et al. 2021). Therefore, this evidence supports the hypothesis that the MBPh-immunoreactive particles were myelin debris-like structures in the normal rat optic nerve.

In the present study, we demonstrate through immunohistochemical analyses: 1) the concentration of MBPh-immunoreactive particles in the distal-most part of the myelinated region; 2) the presence of MBPh-immunoreactive particles containing authentic myelin basic protein; 3) the colocalization of these particles with marker proteins for both neurons and glial cells; 4) the engulfment of these particles by microglia; 5) the morphological similarities between these particles and the MBPh-immunoreactive myelin debris; and 6) the distribution of MBPh-immunoreactive particles in the distal-most part of the myelinated region in both mouse and monkey optic nerves. Finally, based on these observations, we propose the following. Firstly, the concentration of these particles indicates that the distal-most part is subjected to physiologically stressed conditions. Secondly, the concentration of these particles may provide valuable insights into the pathophysiological mechanisms that lead to the localized vulnerability of the myelin sheaths. Lastly, the MBPh-immunoreactive particles and/or the myelin debris could serve as histopathological biomarkers.

## 2 Materials and Methods

### 2.1 Animals and tissue preparation

#### 2.1.1 Normal rats and mice

Male rats (n=13; 12 weeks old; Slc:SD; CLEA Japan, Tokyo, Japan) and male mice (n=3; 8 weeks old; C57BL/6NCrlCrlj; Charles River Laboratories Japan, Yokohama, Kanagawa, Japan) were used in this study. The animals were deeply anesthetized with sodium pentobarbital (50 mg/kg, i.p.), and perfused transcardially with 4 % paraformaldehyde dissolved in 0.1 M sodium phosphate buffer (PB; pH 7.4) at 4°C. The eyeballs, including the optic nerves, were removed from the skull, stored in the same fixative for 48 hours, and then immersed in 30 % saccharose in 0.1 M PB at 4°C until they sank. The eyeballs, along with the optic nerves, were frozen in powdered dry ice, and sectioned in the meridian plane at a thickness of 25 µm on a cryostat. Sections were collected in a cryoprotectant medium (Warr et al. 1981; 33.3% saccharose, 1% polyvinylpyrrolidone (K-30), and 33.3% ethylene glycol in 0.067 M sodium phosphate buffer (pH 7.4) containing 0.067% sodium azide) and stored at –30 °C prior to use (Kawano et al. 2008; Kawano 2015b).

#### 2.1.2 Normal monkeys

The monkey eyeballs, including the optic nerves, were provided by Dr. Shiro Nakagawa (Professor Emeritus, Kagoshima University Graduate School of Medical and Dental Sciences). Male monkeys (n=2; adult; weighing 11.8 to 12.5 kg; *Macaca fuscata*) were initially anesthetized with ketamine hydrochloride (5 mg/kg, i.m.), followed by sodium pentobarbital (40 mg/kg, i.p.). Under deep anesthesia, the monkeys were flushed transcardially with heparinized physiological saline (1,000 units heparin/L), subsequently perfused with 4 % paraformaldehyde dissolved in 0.1 M PB containing 0.2% picric acid at 4°C (Nakagawa, personal communication). The eyeballs, along with the optic nerves, were removed from the skull, stored in 4% paraformaldehyde in 0.1M PB without picric acid for 7 to 9 days, and then immersed in 30 % saccharose in 0.1 M PB at 4°C until they sank. The eyeballs, including the optic nerves, were frozen in powdered dry ice, and sectioned in the meridian plane at a thickness of 40 µm on a freezing microtome. Sections were collected in the cryoprotectant medium and stored at –30 °C prior to use (Kawano 2015a).

#### 2.1.3 Experimental glaucoma model in the rat induced by episcleral vein cauterization (EVC)

Male rats (n=17; 12 weeks old; Slc:SD; CLEA Japan, Tokyo, Japan) were used in this study. All animals were housed in the Kagoshima University animal facility (Kagoshima University, The Center for Advanced Science Research and Promotion, Division of Laboratory Animal Resources and Research) with ad libitum access to food and water under a 12-hour light/12-hour dark cycle at room temperature (23 ± 1 °C) and humidity (55 %). Elevated intraocular pressure was induced by EVC in the left eyes, essentially adhering to a method established by Shareef et al. (1995), which was subsequently followed by Kanamori et al. (2004) and their research group (Naka et al. 2010). The rats were anesthetized through intraperitoneal injection of ketamine hydrochloride (100 mg/kg) and xylazine hydrochloride (10 mg/kg). After making a minimal conjunctival incision, four episcleral veins near the superior, temporal, and inferior rectus muscles were cauterized by diathermy using bipolar cautery forceps. The eyes were flushed with saline and treated with antibiotic ointment (Gentamicin Sulfate ointment; 1mg/g).

After the induction of anesthesia with 2.0% isoflurane in oxygen (2 L/min) delivered to an induction chamber, ketamine hydrochloride (60 mg/kg) and xylazine hydrochloride (6 mg/kg) were administered intraperitoneally. Intraocular pressures (IOPs) were measured in both eyes of anesthetized rats using a rebound tonometer (TonoLab TV02; Icare Finland Oy, Vantaa, Finland) as described by Naka et al. (2010). The device was secured to a ring stand with the probe oriented horizontally. The rats were placed on an adjustable table, and the height of the table was adjusted to ensure that the probe tip was positioned at the center of the cornea at a distance of 2 mm. During each recording session, the tonometer took six measurements, deemed reliable by the internal software, which excluded the highest and lowest readings. Subsequently, the mean IOP was calculated, displayed, and defined as the IOP at the specific time point (Naka et al. 2010).

At 2 weeks and 4 weeks after EVC, the procedures for tissue preparation in the glaucoma rats were the same as those used in the normal rats and mice (Section 2.1.1).

All animal experiments were approved by the Institutional Animal Care and Use Committee of Kagoshima University (rats: MD11112, MD15029, MD18058; mice: MD07068; monkeys: 00205, 00445; glaucoma rats: MD15081, MD18045), and were conducted according to the related guidelines and applicable laws in Japan.

### 2.2 Antibody characterization

Please refer to Table 1 for a list of all primary antibodies used. These antibodies are cataloged in the “Journal of Comparative Neurology antibody database (Version 14)” except for the rabbit anti-glutamine synthetase (GS) antibody.

**TABLE 1.**
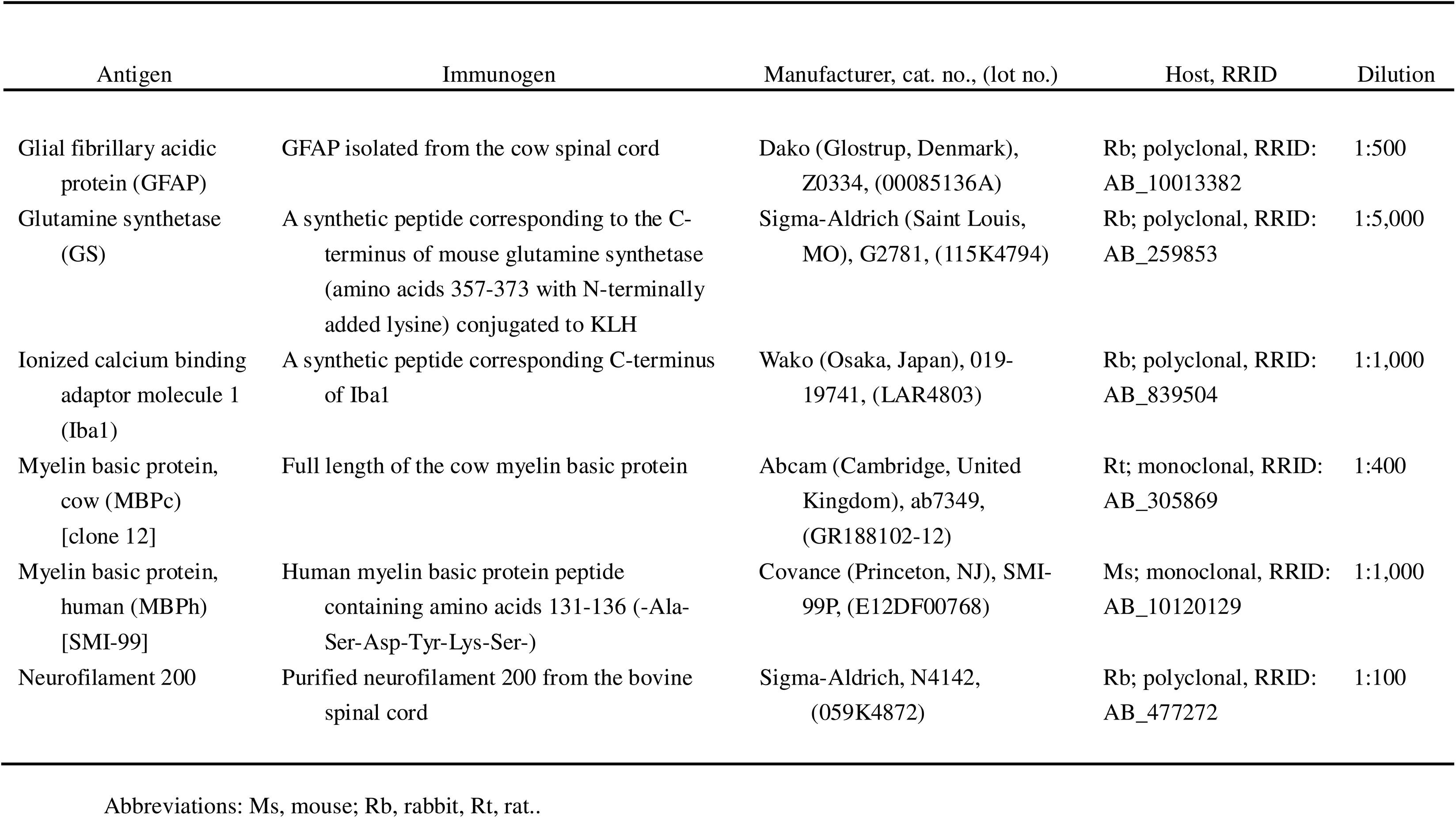
Primary antibodies used in this study.

#### Glial fibrillary acidic protein (GFAP)

The affinity-purified anti-GFAP rabbit antibody (Dako, Glostrup, Denmark) recognizes a single protein band of ≈ 50 kDa in extracts from the mouse retina (Smith et al. 1997; Gaillard et al. 2008). Astrocytes were immunolabeled with this antibody against GFAP in the human optic nerve head (Ye and Hernandez 1995). The staining obtained with this anti-GFAP antibody in the rat was similar to published results on the rat (Saari et al. 1997; Morcos and Chan-Ling 2000; Ju et al. 2005; Chang et al. 2007; Kawano 2015b).

#### Glutamine synthetase (GS)

The rabbit anti-GS antibody (Sigma-Aldrich, Saint Louis, MO, USA) recognizes a single protein band of 45 kDa in extracts from the rat brain. Immunoblotting analysis demonstrates that the staining of GS is specifically inhibited by the GS immunizing peptide (amino acids 357-373 with N-terminally added lysine). This amino acid sequence is identical in human, bovine, rat, hamster, and pig GS, and is highly conserved in chicken GS (single amino acid substitution; manufacturer’s technical information). Müller cells in the retina and glial cells in the optic nerve are labeled with this antibody against GS (Riepe et al. 1977). The staining obtained with this antibody in the rat was similar to that previously reported in the mouse (Haverkamp and Wässle 2000; Hojo et al. 2000; Kawano et al. 2008) and the rat (Riepe et al. 1977; Zabouri et al. 2011; Kawano 2015b).

#### Ionized calcium binding adaptor molecule 1 (Iba1)

Microglia and macrophages are immunostained using the rabbit polyclonal antibody against Iba1 (Wako, Osaka, Japan); however, neurons and astrocytes do not exhibit immunostaining with this antibody (Ito et al. 2001; manufacturer’s technical information). The Iba1 antibody recognizes a single protein band of 17 kDa corresponding to the Iba1 protein in extracts from rat brain microglia cultures and from several human monocytic cell lines (Imai et al. 1996). This antibody stains retinal and optic nerve microglia in both the mouse (Bosco et al. 2008; Santos et al. 2008; Bosco et al. 2011) and in the rat (Naskar et al. 2002; Johnson et al. 2007; Zhang et al. 2009). The staining obtained with this antibody against Iba1 was similar to that previously reported in the rat (Zhang et al. 2009).

#### Myelin basic protein, cow (MBPc)

The rat monoclonal antibody against the full-length cow myelin basic protein (clone 12; Abcam, Cambridge, United Kingdom) recognizes two protein bands of 19 and 26 kDa on immunoblots of mouse brain tissue lysate (manufacturer’s technical information). This antibody stains myelin sheaths of cultured oligodendrocyte precursor cells obtained from the postnatal day 7 rat brain (Dugas et al. 2006). The staining obtained with this antibody against cow myelin basic protein was similar to that previously reported in the rat (Morcos and Chan-Ling 2000).

#### Myelin basic protein, human (MBPh)

The mouse monoclonal antibody against human myelin basic protein (clone SMI-99; Covance) detects four bands between 14 and 21 kDa, corresponding to four myelin basic protein (MBP) isoforms on immunoblots of the mouse cerebellum (Dyer et al. 1996; Talos et al. 2006). The SMI-99 antibody detects MBP from most mammalian species. The pig and chicken MBP do not react, while the guinea pig MBP has slight reactivity. This antibody does not react with the 14kDa form of the rat MBP. The SMI-99 antibody detects the developing and adult myelin, and distinguishes oligodendrocytes from astrocytes, microglia, neurons and other cells in brain sections (manufacturer’s technical information). The staining obtained with this antibody against human myelin basic protein was similar to that previously reported in the rat (Morcos and Chan-Ling 2000), except for the concentration of the MBPh-immunoreactive particles.

#### Neurofilament 200-kDa heavy chain (neurofilament 200; NF200)

The polyclonal anti-NF200 antibody (Sigma-Aldrich, Saint Louis, MO, USA) recognizes a single protein band of 200 kDa in extracts from the rat brain cytosolic S1 fraction (manufacturer’s technical information) and in those from the mouse brain (Benvegnù et al. 2010). The antibody shows wide species cross-reactivity (manufacturer’s technical information). The staining obtained with this antibody against NF200 was similar to published results on the mouse (Howell et al. 2007) and the rat (Naka et al. 2010; Kawano 2015b).

### 2.3 Immunohistochemistry

Sections were processed using double-label immunohistochemistry as previously described (Kawano et al. 2008; Kawano 2015b), except for the detection of MBP by using the rat monoclonal anti-MBPc antibody. Free-floating sections were pre-incubated for 2 hours with a 10% normal goat serum (NGS) blocking solution at 4 °C and then immunoreacted for 4 days with a mixture of rabbit and mouse primary antibodies in a 10% NGS blocking solution at 4 °C (Table 1). After two rinses for 10 minutes in 0.02M phosphate buffered saline (PBS) containing 0.3% Triton X-100 (PBST), the sections were incubated with a mixture of two secondary antibodies in PBS containing 5% NGS and 0.3% Triton X-100 for 24 hours at 4 °C. The two secondary antibodies used were Alexa Fluor 488 conjugated with the F(ab’)2 fragment of goat anti-rabbit IgG (H+L) (1:200; Molecular Probes, Eugene, OR, USA) and Alexa Fluor 594 conjugated to the F(ab’)2 fragment of goat anti-mouse IgG (H+L) (1:200; Molecular Probes). The sections were washed once for 10 minutes in PBST and then twice in PBS. The sections were mounted onto hydrophilic silanized slides (Dako Japan, Tokyo, Japan) by using an equal-parts mixture of a 0.6% gelatin solution and PBS. After being air-dried, the sections were subjected to nuclear staining by using a bisBenzimide (bBM; Hoechst 33258, Sigma-Aldrich; 0.1 mg/ml) solution and coverslipped with VECTASHIELD mounting medium (Vector Laboratories, Burlingame, CA, USA).

In instances of double-label immunohistochemistry by using the rabbit polyclonal anti-GS antibody and the rat monoclonal anti-MBPc antibody as primary antibodies, Alexa Fluor 488 conjugated with the F(ab’)2 fragment of goat anti-rabbit IgG (H+L) (1:500; Molecular Probes) and Alexa Fluor 594 conjugated to goat anti-rat IgG (H+L) preadsorbed (1:200; Abcam) were employed as secondary antibodies.

In instances of double-label immunohistochemistry by using the rat monoclonal anti-MBPc antibody and the mouse monoclonal anti-MBPh antibody as primary antibodies, Alexa Fluor 488 conjugated to goat anti-rat IgG (H+L) cross-adsorbed (1:500; Thermo Fisher Scientific, Waltham, MA, USA) and Alexa Fluor 594 conjugated to goat anti-mouse IgG (H+L) highly cross-adsorbed (1:500; Thermo Fisher Scientific) were employed as secondary antibodies. Due to the slight immunoreaction of Alexa Fluor 594 conjugated to goat anti-mouse IgG (H+L) highly cross-adsorbed with the rat monoclonal anti-MBPc antibody, the primary antibodies were applied serially rather than simultaneously as follows. After 2 hours preincubation with the 10% NGS blocking solution, free-floating sections were incubated overnight with the mouse monoclonal anti-MBPh antibody. Following two rinses in PBST, the sections were immunoreacted for 3 hours with Alexa Fluor 594 conjugated to goat anti-mouse IgG (H+L) highly cross-adsorbed. After two washes in PBST, the sections were incubated overnight with the rat monoclonal anti-MBPc antibody. After two rinses in PBST, the sections were immunoreacted for 3 hours with Alexa Fluor 488 conjugated to goat anti-rat IgG (H+L) cross-adsorbed. The subsequent procedures were the same as described above.

To eliminate the possibility of any cross-reaction between the secondary and primary antibodies from the different species, one of the two primary antibodies was removed. No cross-reactivity was observed in these control experiments (Supplementary Figure 1).

In normal rat cases, each staining protocol was conducted on a minimum of 3 optic nerves from 3 separate rats. GS/MBPh staining^1^, MBPc/MBPh staining, and Iba1/MBPh staining protocols were executed on 5 optic nerves from 5 separate rats each. GS/MBPc staining, MBPc control staining^2^, MBPh control staining, NF200/MBPh staining, and GFAP/MBPh staining protocols were performed on 3 optic nerves from 3 separate rats each. In glaucoma rat cases, GS/MBPh staining protocol was conducted on a total of 11 optic nerves in the operated side and a total of 11 optic nerves in the non-operated side from 11 separate rats. In normal mouse cases, GS/MBPh staining protocol was executed on a total of 3 optic nerves from 3 separate mice. In normal monkey cases, GS/MBPh staining protocol was performed on a total of 2 optic nerves from 2 separate monkeys.

### 2.4 Photomicrographs

Fluorescent photomicrographs were obtained using either LSM700 or LSM900 confocal laser scanning microscopes (Carl Zeiss Jena GmbH, Jena, Germany) at the Joint Research Laboratory, Kagoshima University Graduate School of Medical and Dental Sciences (see Figure Legends). The images were transferred to Adobe Photoshop CS5 (Adobe Systems, San Jose, CA, USA), where brightness and contrast were adjusted. No other adjustments were made.

### 2.5 Image analysis

The quantitation of all images was performed using ImageJ2 (Version 2.9.0/1.53t; developed by Wayne Rasband, National Institute of Mental Health, Bethesda, MD, USA).

### 2.6 Quantitative morphological analyses comparing MBPh-immunoreactive particles and myelin debris

#### 2.6.1 Measurement of the mean area, mean perimeter, and of mean shape descriptors of MBPh-immunoreactive particles

Area, perimeter, and shape descriptors of each MBPh-immunoreactive particle were measured in a trapezoid-like image, which showed the distribution of the particles in the distal-most part of the myelinated region. The bottom of the image was set at the border between the unmyelinated and myelinated regions. The longitudinal length of the image was 250 µm. To exclude the background from the measurement area, the boundaries of the myelinated regions were cut along the pia mater.

For each image, the process conducted by the ImageJ2 program included the following: (a) executing the Subtract Background function to remove smooth continuous background from the image (Rolling Ball Algorithm; Radius: 10.0 pixels); (b) performing the Median filter function to reduce noise in the image (Radius: 2.0 pixels); (c) conducting the Mean filter function to smooth the image (Radius: 2.0 pixels); (d) establishing the threshold in order to binarize the image (Auto; Moments); (e) applying the watershed; (f) analyzing particles to obtain area, perimeter, and shape descriptors (Circularity; AR (aspect ratio); Roundness; Solidity) of each MBPh-immunoreactive particle, and to count MBPh-immunoreactive particles (size, 1.0 µm^2^-infinity; circularity, 0.00-1.00; check “Clear results”, “Add to manager”, “Exclude on edges”, and “Include holes”).

Mean area, mean perimeter, and mean shape descriptors of MBPh-immunoreactive particles were calculated manually based on the outputs of the measurement processes described above.

#### 2.6.2 Measurement of the density of MBPh-immunoreactive particles

The number of MBPh-immunoreactive particles were counted as described above (Section 2.6.1). The area of the trapezoid-like image, in which the particles were distributed, was measured as follows. For each image, the process performed by the ImageJ2 program included the following: (a) setting the threshold 1-255 to binarize the image (Manual). (b) analyzing particles to measure area (size, 100 µm^2^-infinity; circularity, 0.00-1.00; check “Clear results”, “Add to manager”, and “Include holes”). The density of MBPh-immunoreactive particles was manually calculated based on the number of particles and on the area in which the particles were distributed.

### 2.7 Statistical analysis

No statistical methods were used to predetermine group sample size. However, our group sample sizes were similar to those in previously published studies by our group and others (Melo et al. 2006; Balaratnasingam et al. 2009; Kawano 2015b). All biological replicates (n) were derived from at least three independent experiments. Unless otherwise specified, no data were excluded from analysis. Individual experiment values are represented in bar graphs or box and whisker plots. In the bar graphs, the edges of the bars indicate the means, and the end of the whiskers represents the standard deviation of the distribution. In the box and whisker plots, the upper whisker indicates the maximum value while the lower one represents the minimum value. The thick horizontal line in the box shows the median. The upper and lower borders of the box indicate the interquartile range.

All statistical analyses were conducted by using EZR (Easy R; Version 1.55; Saitama Medical Center, Jichi Medical University, Saitama, Japan), which serves as a graphical user interface for R (Version 4.1.3; The R Foundation for Statistical Computing, Vienna, Austria). More specifically, it is a modified version of R Commander (Version 2.7-2; Fox 2017) that is designed to add statistical functions frequently used in biostatistics (Kanda 2013). Data were evaluated using the Kolmogorov-Smirnov test for normal distribution, the Shapiro-Wilk normality test, the two-variances F-test for homogeneity of variance, the two-sample t-test, and the Mann-Whitney U test. A p-value of < 0.05 was considered statistically significant.

## 3 Results

### 3.1 Distribution of MBPh-immunoreactive particles in the normal rat optic nerve

MBPh-immunoreactive particles were concentrated in the distal-most part of the myelinated region in the normal rat optic nerve (Figure 1A). However, a small number of MBPc-immunoreactive particles were detected in a similar part of the myelinated region (Figure 1B). Interestingly, the majority of both MBPh-immunoreactive and MBPc-immunoreactive particles were distributed on MBP-immunoreactive myelinated nerve fibers (Figure 1A, B).

### 3.2 The anti-MBPh antibody demonstrated immunoreactivity with the authentic myelin basic protein, and the MBPh-immunoreactive particles were found to contain the authentic myelin basic protein

Fluorescent double immunohistochemistry by using the anti-MBPh and anti-MBPc antibodies demonstrates that the majority of MBPh-immunoreactive particles were also immunolabeled with the anti-MBPc antibody, which targets a different amino acid sequence from that of the anti-MBPh antibody (Figure 2D). These facts indicate that the anti-MBPh antibody is immunoreactive for the authentic myelin basic protein and that the MBPh-immunoreactive particles contain the authentic myelin basic protein (Figure 2D).

**FIGURE 2.**
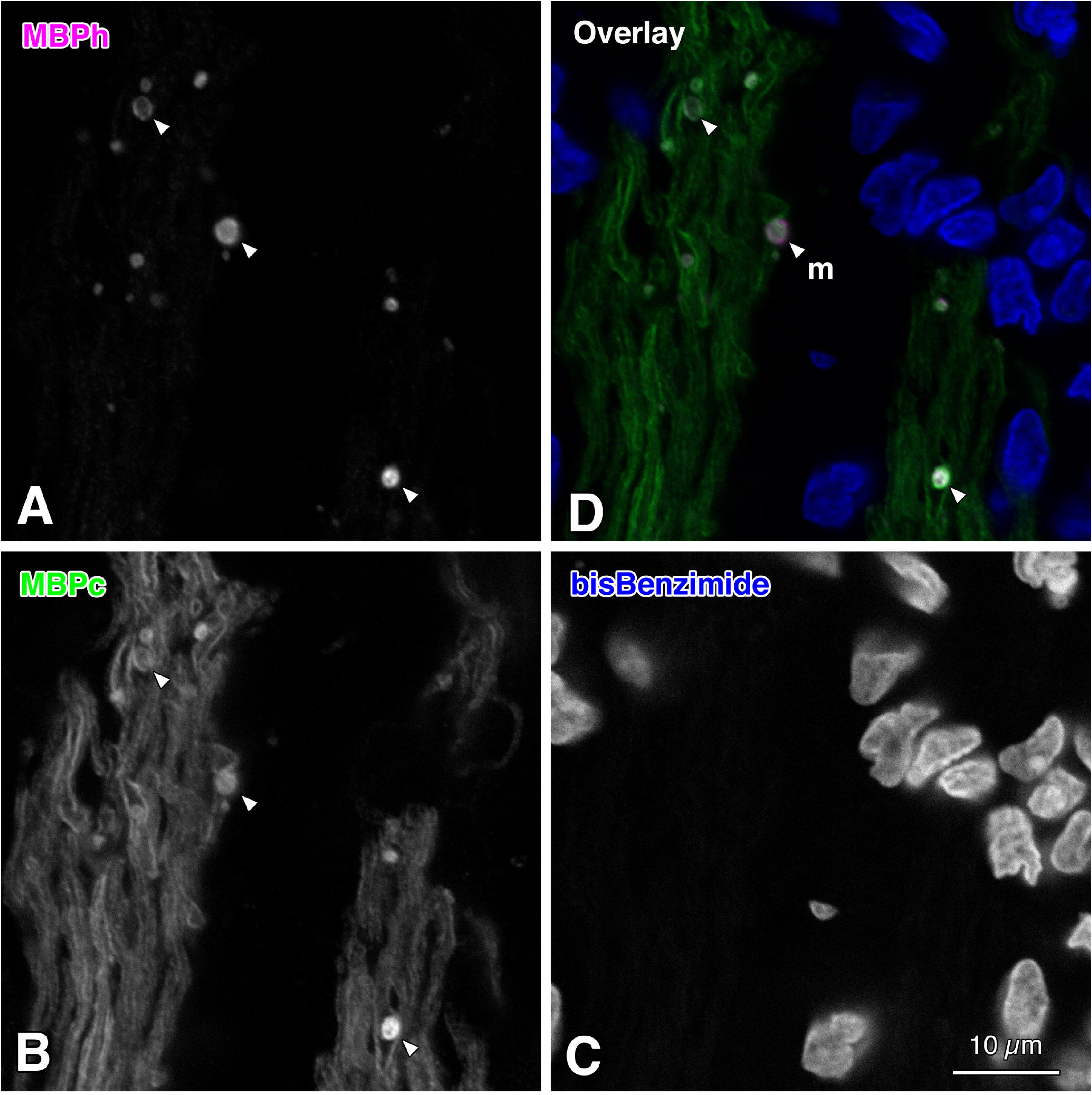
Fluorescent double immunohistochemistry by using two monoclonal anti-myelin basic protein (MBP) antibodies that target different amino acid sequences. Images show a small square region of a longitudinal section through the paramedian part in the distal (anterior)-most part of the myelinated region in the normal rat optic nerve. **A:** Particles were labeled by using a mouse monoclonal anti-human myelin basic protein (MBPh) antibody (clone SMI-99; Covance, Princeton, NJ, USA; Alexa Fluor 594 label; magenta in **D**). This antibody reacts with Ala-Ser-Asp-Tyr-Lys-Ser (ASDYKS) in position 131-136 of the classic human myelin basic protein. **B:** Particles and myelinated nerve fibers were labeled with a rat monoclonal anti-cow myelin basic protein (MBPc) antibody (clone 12; Abcam, Cambridge, United Kingdom; Alexa Fluor 488 label; green in **D**). This antibody reacts with Asp-Glu-Asn-Pro-Val-Val (DENPVV) in position 82-87 of the full-length cow myelin basic protein. **C:** Cell nuclei labeled with bisBenzimide (Hoechst 33258; blue in **D**). **D:** A color overlay image of panels **A-C**. The arrowheads in panels **A**, **B**, and **D** indicate MBPh-immunoreactive particles that are also immunolabeled with the anti-MBPc antibody. These images were taken with an LSM 700 confocal microscope (Carl Zeiss, Jena, Germany). Note that the majority of MBPh-immunoreactive particles are also immunolabeled with the anti-MBPc antibody, which targets an amino acid sequence different from that of the anti-MBPh antibody (**D**). These facts demonstrate that the anti-MBPh antibody was immunoreactive for the authentic myelin basic protein and that the MBPh-immunoreactive particles contained the authentic myelin basic protein. Additionally, the majority of the MBP-immunoreactive particles are distributed on MBPc-immunoreactive myelinated nerve fibers (**D**). m, myelinated region. Scale bar = 10 µm in the lower right corner of **C** for both **A-B** and **D**.

The control studies show that cross-immunoreactions were not detectable between the mouse monoclonal anti-MBPh antibody and the Alexa Fluor 488 conjugated goat anti-rat secondary antibody (Supplementary Figure 1C), as well as between the rat monoclonal anti-MBPc antibody and the Alexa Fluor 594 conjugated goat anti-mouse secondary antibody (Supplementary Figure 1J). These findings indicate that the fluorescent double immunohistochemistry by using the mouse monoclonal anti-MBPh antibody and the rat monoclonal anti-MBPc antibody was genuine rather than artificial. Consequently, the MBPh and MBPc double-immunoreactive particles seen in Figure 2D were not false positives but authentic.

### 3.3 Colocalization of neural and glial cell marker proteins in the MBPh-immunoreactive particles

#### 3.3.1 Colocalization of GFAP (glial fibrillary acidic protein) in the MBPh-immunoreactive particles

Moderate GFAP-immunoreactivity was observed in the MBPh-immunoreactive particles (see GFAP-immunoreactivity in the particles indicated by the arrowheads in Figure 3A, 3B, and 3D).

**FIGURE 3.**
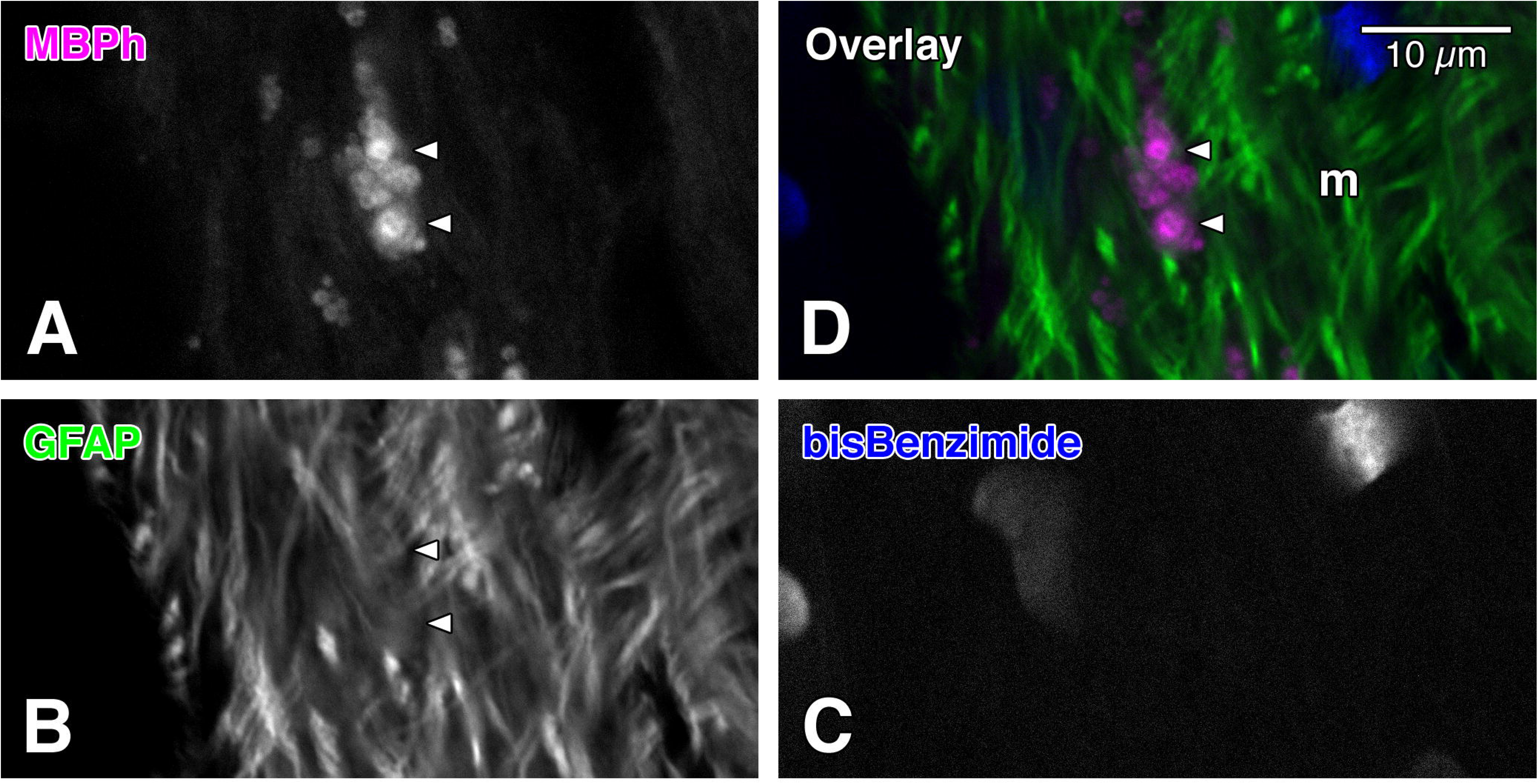
Images showing the distribution of glial fibrillary acidic protein (GFAP) and of myelin basic protein-immunoreactive particles visualized by using fluorescent double immunohistochemistry. The images represent a small rectangular region of a longitudinal section through the paramedian part in the distal (anterior)-most part of the myelinated region in the normal rat optic nerve. Several particles strongly labeled in white (**A**) and in magenta (D) were labeled with a mouse monoclonal anti-human myelin basic protein (MBPh) antibody (clone SMI-99; Covance, Princeton, NJ, USA; Alexa Fluor 594 label). GFAP was labeled with an anti-GFAP antibody (Dako, Glostrup, Denmark; Alexa Fluor 488 label; **B**; green in **D**). Cell nuclei were labeled with bisBenzimide (Hoechst 33258; **C**; blue in **D**). The arrowheads indicate strongly MBPh-immunoreactive particles. These images were taken with an LSM 700 confocal microscope (Carl Zeiss, Jena, Germany). Note that moderate GFAP-immunoreactivity is observed in the MBPh-immunoreactive particles (see GFAP-immunoreactivity pointed by the arrowheads in **B**). m, myelinated region. Scale bar = 10 µm in the upper right corner of **D** for **A-C**.

#### 3.3.2 Colocalization of GS (glutamine synthetase) in the MBPh-immunoreactive particles

GS-immunoreactivity was clearly observed in the MBPh-immunoreactive particles (see GS-immunoreactivity in the particles indicated by the arrowheads in Figure 4A, 4B, and 4D).

**FIGURE 4.**
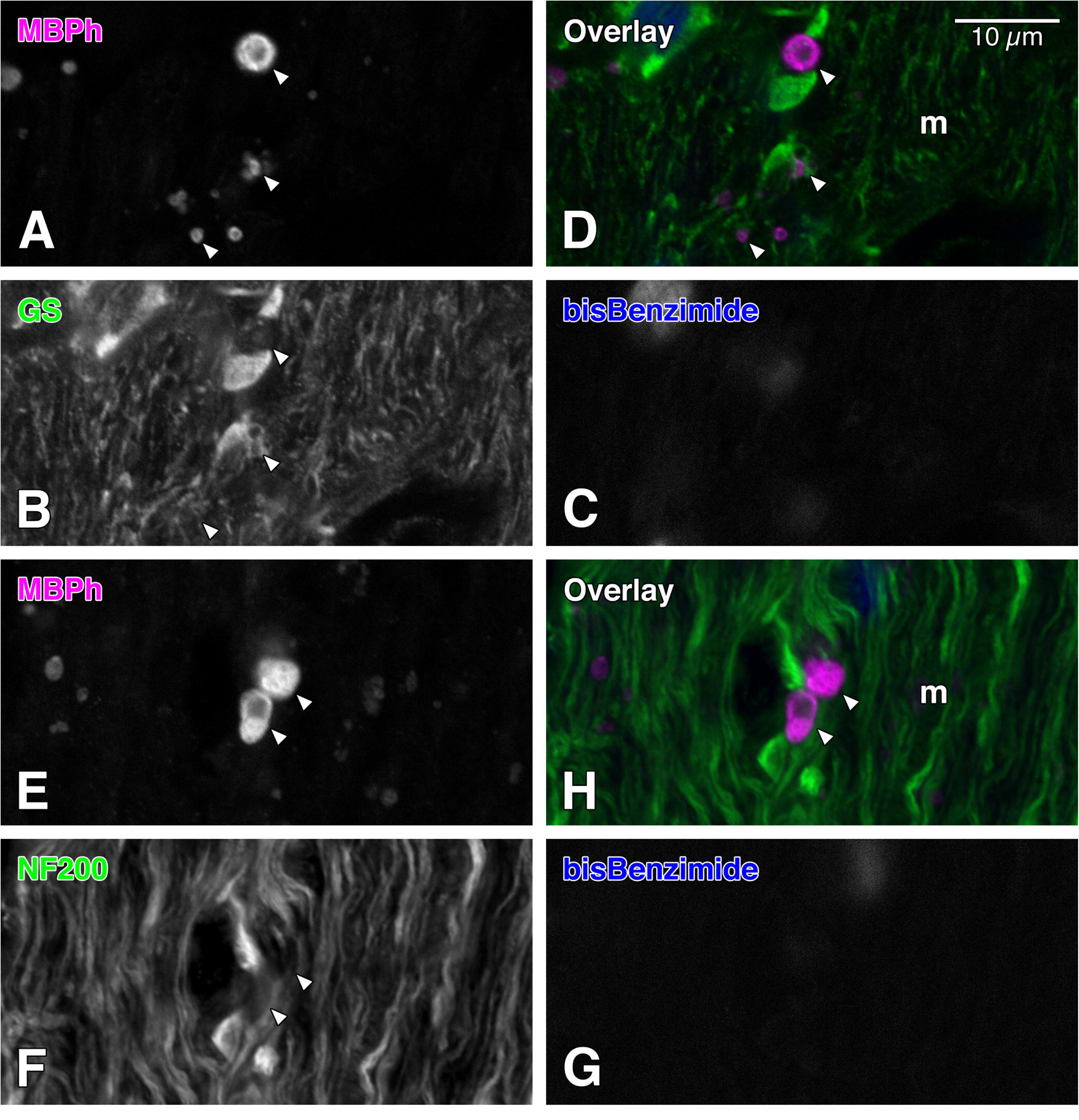
A-D: Images showing the distribution of glutamine synthetase (GS) and of myelin basic protein-immunoreactive particles visualized by using fluorescent double immunohistochemistry. The images represent a small rectangular region of a longitudinal section through the paramedian part in the distal (anterior)-most part of the myelinated region in the normal rat optic nerve. Several particles colored in white (**A**) and in magenta (**D**) were labeled with a mouse monoclonal anti-human myelin basic protein (MBPh) antibody (clone SMI-99; Covance, Princeton, NJ, USA; Alexa Fluor 594 label). The GS protein was labeled with an anti-GS antibody (Sigma-Aldrich, Saint Louis, MO, USA; Alexa Fluor 488 label; **B**; green in **D**). Cell nuclei were labeled with bisBenzimide (Hoechst 33258; **C**; blue in **D**). The arrowheads indicate strongly-to-moderately MBPh-immunoreactive particles (**A-B**, **D**). These images were taken with an LSM 700 confocal microscope (Carl Zeiss, Jena, Germany). Note that moderate GS-immunoreactivity is observed in the MBPh-immunoreactive particles (see GS-immunoreactivity pointed by the arrowheads in **B**). **E-H:** Images showing the distribution of the neurofilament 200-kDa heavy chain (NF200) protein and of MBPh-immunoreactive particles visualized by using fluorescent double immunohistochemistry. The images represent a small rectangular region of a longitudinal section through the paramedian part in the distal-most part of the myelinated region in the normal rat optic nerve. Several particles labeled in white (**E**) and in magenta (**H**) were labeled with the anti-MBPh antibody (Alexa Fluor 594 label). The NF200 protein was labeled with an anti-NF200 antibody (Sigma-Aldrich; Alexa Fluor 488 label; **F**; green in **H**). Cell nuclei were labeled with bisBenzimide (Hoechst 33258; **G**; blue in **H**). The arrowheads indicate strongly MBPh-immunoreactive particles. These images were taken with the LSM 700 confocal microscope (Carl Zeiss). Note that weak-to-moderate NF200-immunoreactivity is observed in the MBPh-immunoreactive particles (see NF-200 immunoreactivity pointed by the arrowheads in **F**). m, myelinated region. Scale bar = 10 µm in the upper right corner of **D** for **A-C**, and for **E-H**.

#### 3.3.3 Colocalization of NF200 (neurofilament 200-kDa heavy chain; neurofilament 200) in the MBPh-immunoreactive particles

Moderately NF200-immunoreactive fibers were observed in the MBP-immunoreactive particles (see NF200-immunoreactive fibers in the particles indicated by the arrowheads in Figure 4E, 4F, and 4H).

#### 3.3.4 Distribution of Iba1 (ionized calcium binding adapter molecule 1)-labeled microglia, and of the MBPh-immunoreactive particles

The majority of MBPh-immunoreactive particles were isolated from Iba1-labeled microglia (Figure 5D). It was observed that at least five MBPh-immunoreactive particles had been engulfed by an Iba1-labeled microglial cell (Figure 5D, inset).

**FIGURE 5.**
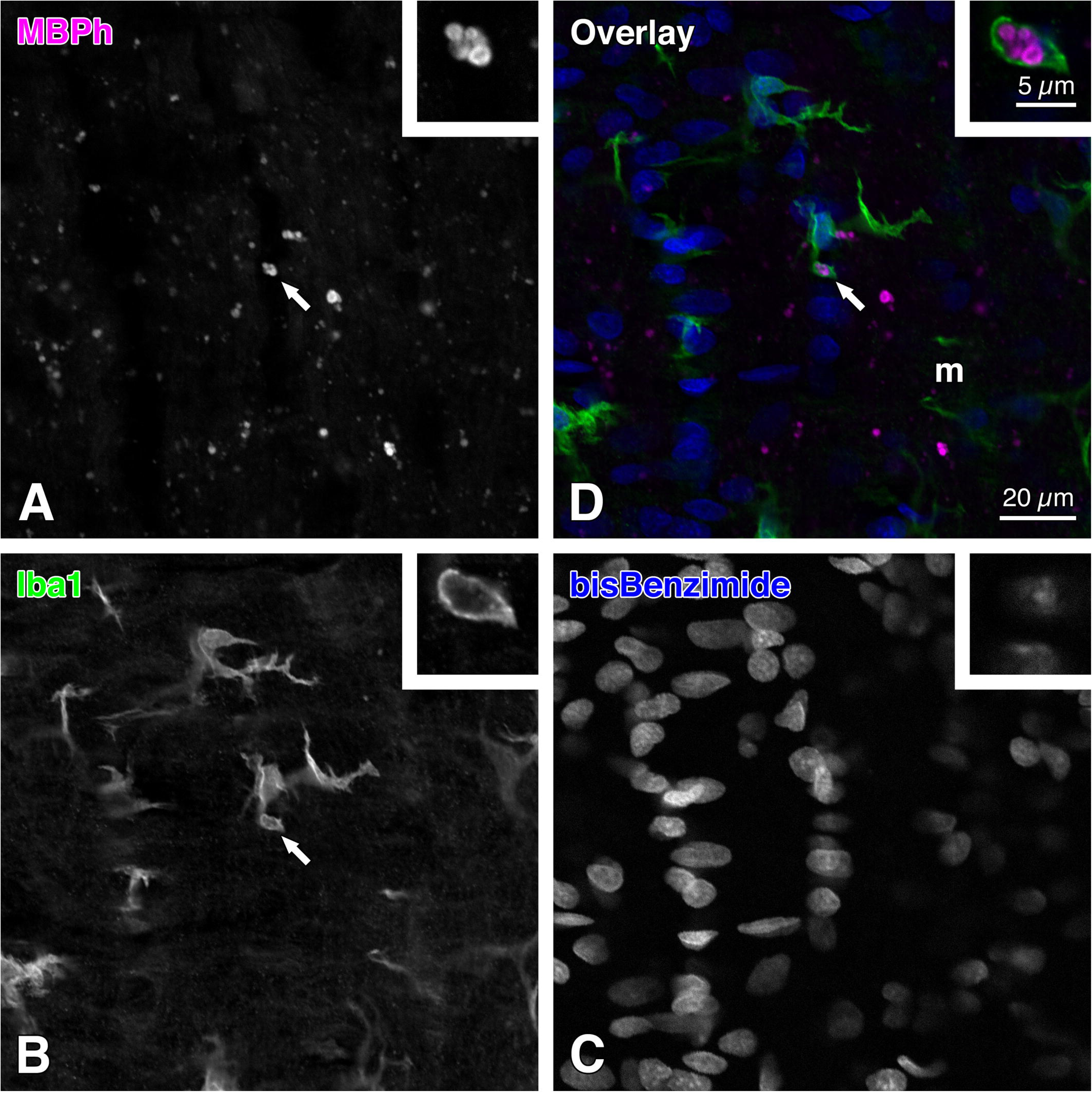
Images showing the distribution of ionized calcium binding adapter molecule 1 (Iba1)-labeled microglia and of myelin basic protein-immunoreactive particles visualized by using fluorescent double immunohistochemistry. These images represent a small square region of a longitudinal section through the paramedian part in the distal (anterior)-most part of the myelinated region in the normal rat optic nerve. **A:** Particles were labeled with a mouse monoclonal anti-human myelin basic protein (MBPh) antibody (clone SMI-99; Covance, Princeton, NJ, USA; Alexa Fluor 594 label; magenta in **D**). **B:** Microglia were labeled with a rabbit polyclonal anti-Iba1 antibody (Wako Pure Chemical Industries, Osaka, Japan; Alexa Fluor 488 label; green in **D**). **C:** Cell nuclei were labeled with bisBenzimide (Hoechst 33258; blue in **D**). **D:** A color overlay image of panels **A-C**. The arrows in panels **A**, **B**, and **D** indicate MBPh-immunoreactive particles engulfed by an Iba1-labeled microglial cell. **Insets** provide higher-magnification photomicrographs of these particles and the Iba1-labeled microglial cell. These images were taken with an LSM 700 (**A-D**) or an LSM 900 (**insets**) confocal microscope (Carl Zeiss, Jena, Germany). Note that the majority of MBPh-immunoreactive particles are isolated from Iba1-labeled microglia (**D**). Additionally, at least five MBPh-immunoreactive particles are engulfed by the Iba1-labeled microglial cell (**insets** of **A**, **B**, and **D**). m, myelinated region. Scale bar = 20 µm in the lower right corner of **D** for **A-C**; 5 µm in the lower right corner of the **inset** of **D** for the **insets** of **A-C**.

### 3.4 A neuroanatomical comparison between the MBPh-immunoreactive particles and the MBPh-immunoreactive myelin debris

The dimensions of MBPh-immunoreactive particles (Figures 6C-D, 7A-E) were generally comparable to those of MBPh-immunoreactive myelin debris (Figures 6A-B, 7F-H).

**FIGURE 6.**
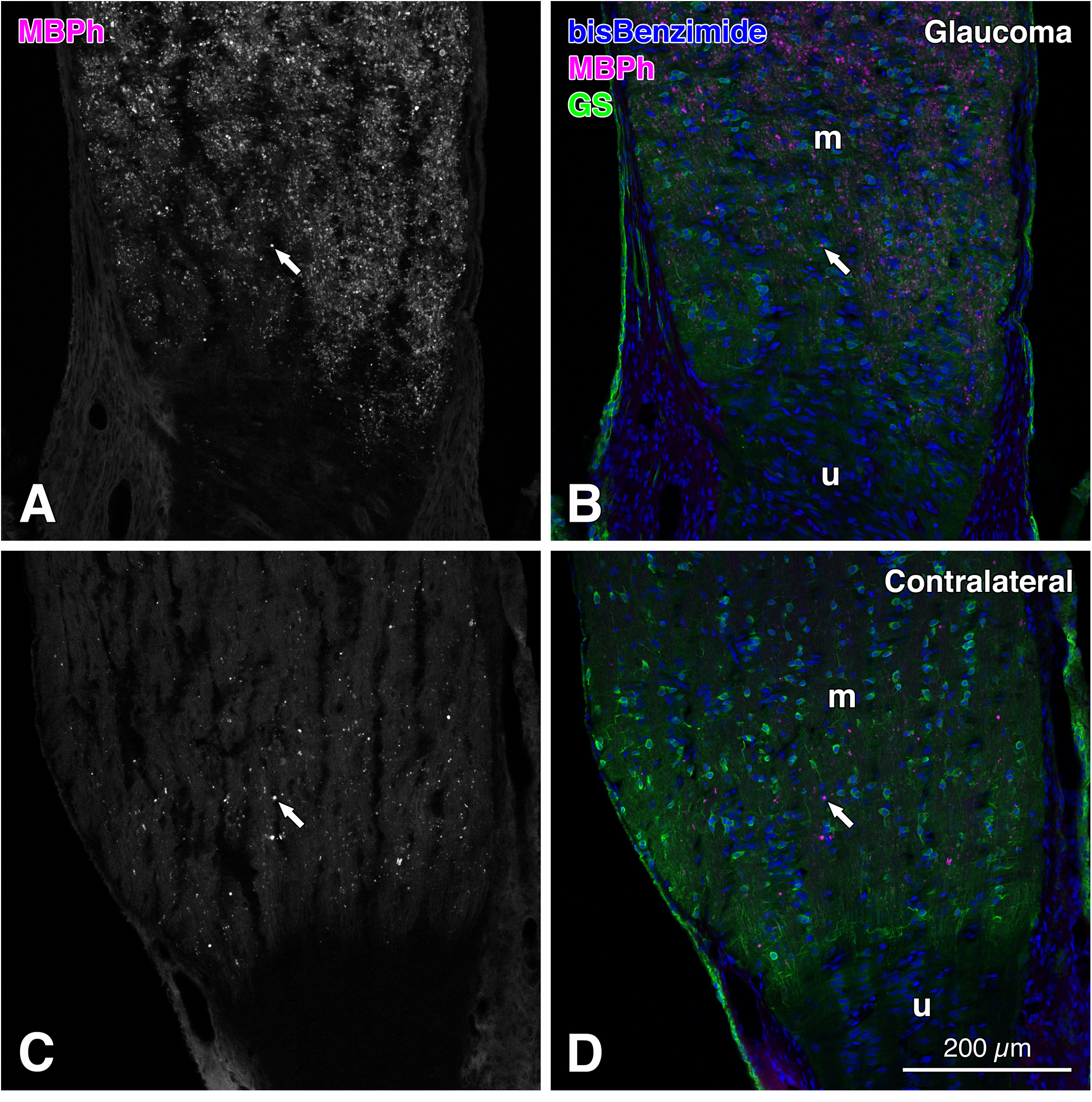
Images showing the distribution of myelin basic protein-immunoreactive myelin debris and of glutamine synthetase (GS) in the glaucoma rat optic nerve visualized by using fluorescent double immunohistochemistry (case code: glaucoma rat 3). Panels **A** and **B** demonstrate the distribution of the myelin debris in the glaucomatous (left) optic nerve. Panels **C** and **D** represent myelin basic protein-immunoreactive particles in the contralateral (right) optic nerve. The images were taken from longitudinal sections through the paramedian part in the distal (anterior)-most part of the myelinated region in the optic nerves. Cell nuclei were labeled with bisBenzimide (Hoechst 33258; blue in **B**, **D**). The GS protein was labeled with an anti-GS antibody (Sigma-Aldrich, Saint Louis, MO, USA; Alexa Fluor 488 label; green in **B**, **D**). Particles colored in white (**A**, **C**) and in magenta (**B**, **D**) were labeled with a mouse monoclonal anti-human myelin basic protein (MBPh) antibody (clone SMI-99; Covance, Princeton, NJ, USA; Alexa Fluor 594 label). The arrows indicate MBPh-immunoreactive myelin debris in the glaucomatous optic nerve (**A**, **B**) and MBPh-immunoreactive particles in the contralateral optic nerve (**C**, **D**). These images were taken with an LSM 700 confocal microscope (Carl Zeiss, Jena, Germany). Note that the size of the MBPh-immunoreactive particles (**C**, **D**) is broadly similar to that of MBPh-immunoreactive myelin debris (**A**, **B**). m, myelinated region; u, unmyelinated region. Scale bar = 200 µm in the lower right corner of **D** for **A-C**.

We next quantitatively compared the mean sizes (area and perimeter) and mean shape descriptors (circularity, AR (aspect ratio), roundness, and solidity) of MBPh-immunoreactive particles in the normal rat (NR) optic nerve with those of MBPh-immunoreactive myelin debris in the damaged optic nerve of the glaucoma rat (GR; Table 2). The ratios of NR means to GR means ranged from 0.971 to 1.029. Therefore, all observed differences in sizes and shape descriptors between the particles and the myelin debris were less than 3 percent.

**TABLE 2.**
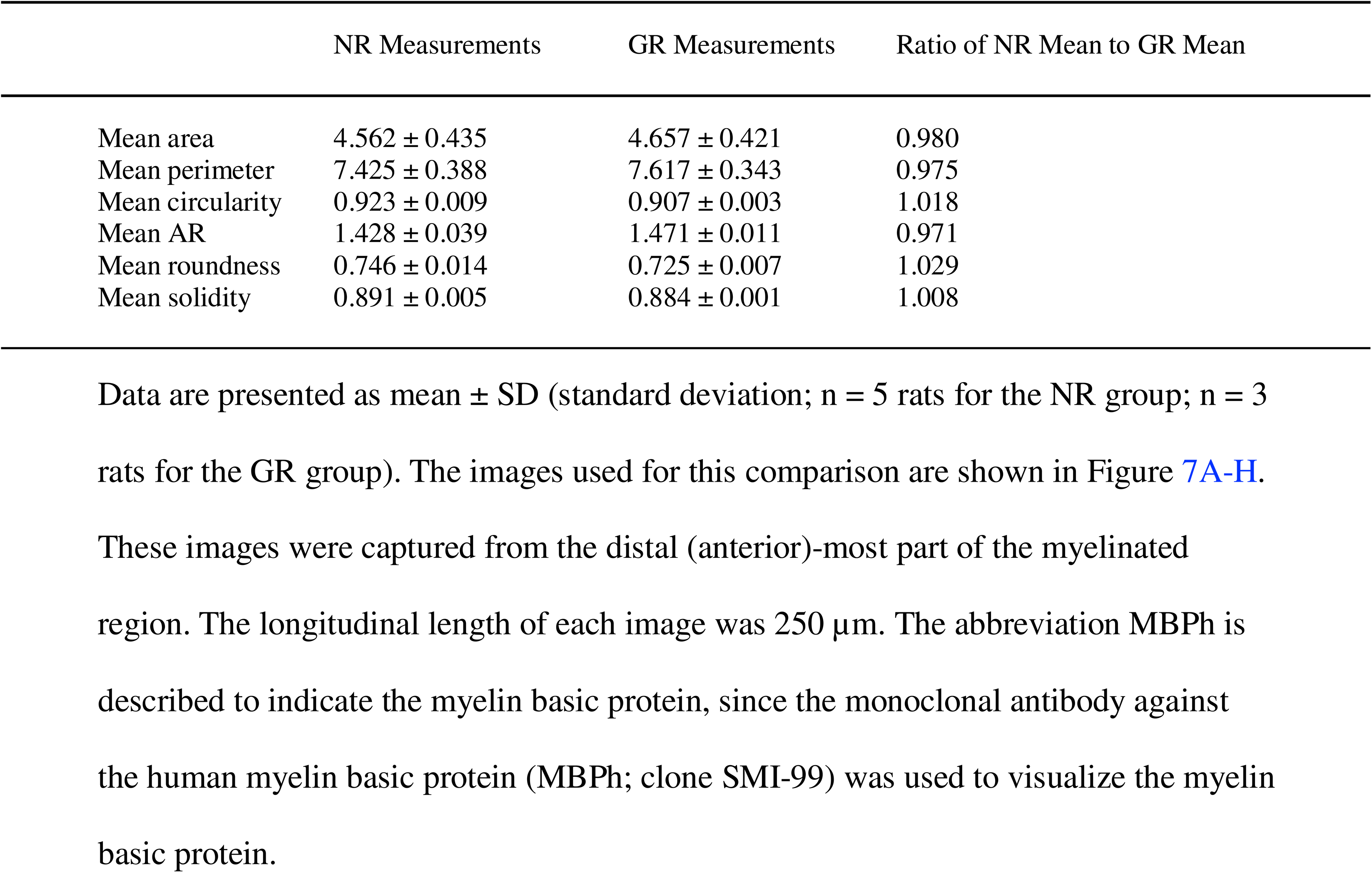
Comparison of the mean sizes (area (µm^2^) and perimeter (µm)) and mean shape descriptors (circularity, AR (aspect ratio), roundness, and solidity) of MBPh-immunoreactive particles in the normal rat (NR) optic nerve with those of MBPh-immunoreactive myelin debris in the damaged optic nerve of the glaucoma rat (GR).

The results of two-sample t-tests (p < 0.05; n = 5 in the normal rat, n=3 in the glaucoma rat) were as follows: mean area, p = 0.773; mean perimeter, p = 0.508; mean circularity, p = 0.032; mean AR (aspect ratio), p = 0.115. The results of Mann-Whitney *U* tests (p < 0.05; n = 5 in the normal rat, n=3 in the glaucoma rat) were as follows: mean roundness, p = 0.036; mean solidity, p = 0.134. Thus, the p-values of both mean circulatory and mean roundness were less than 0.05. Accordingly, the mean circularity and mean roundness of the MBPh-immunoreactive particles were significantly different from those of the MBPh-immunoreactive myelin debris. Therefore, only 2 out of 6 differences in the mean sizes and the mean shape descriptors between the particles and the myelin debris were statistically significant (Figure 8).

**FIGURE 7.**
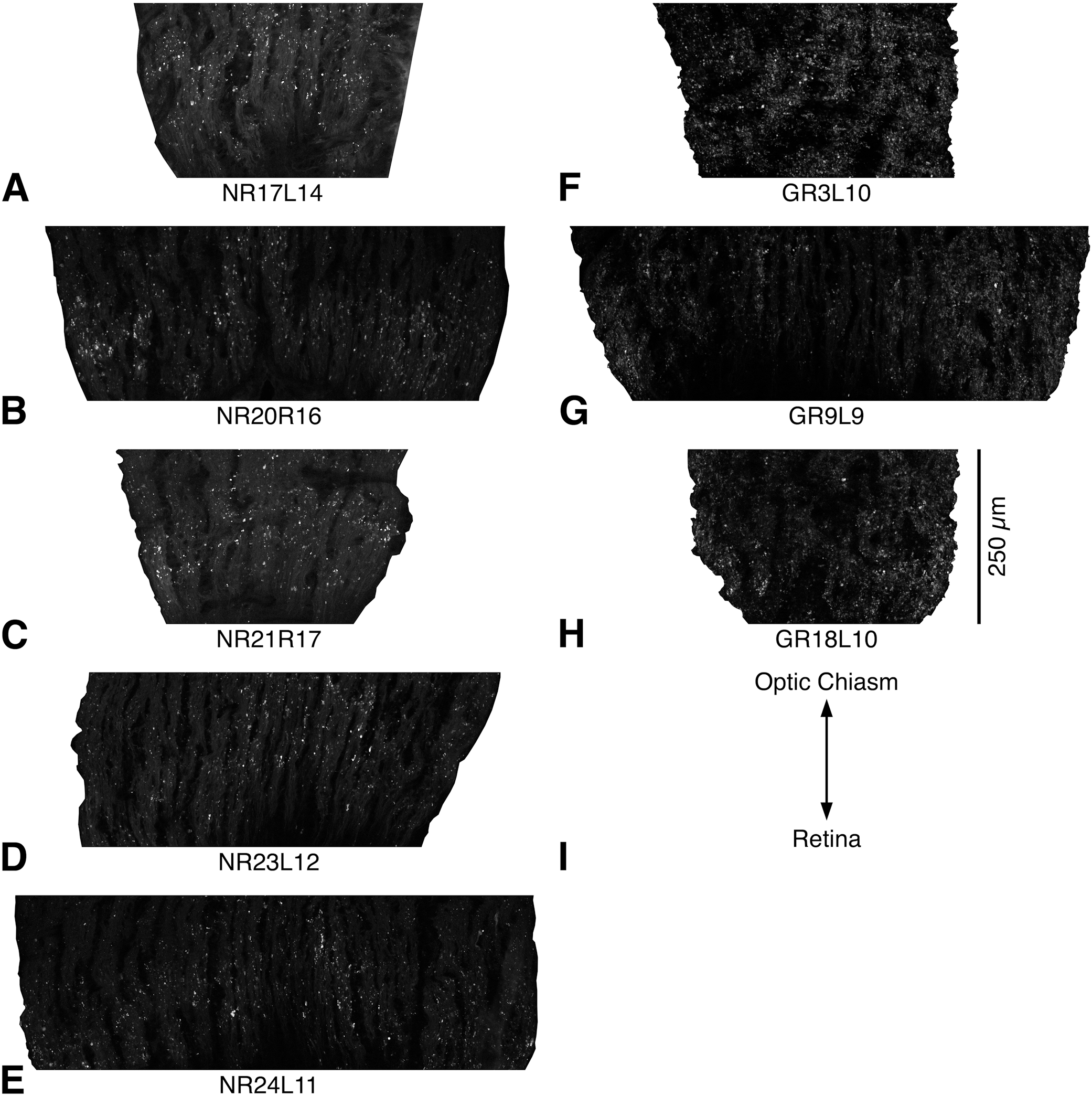
A-H: Images used for statistical analyses of differences in sizes and in shape descriptors between myelin basic protein (MBP)-immunoreactive particles in the distal (anterior)-most part of the myelinated region in the normal rat optic nerve (**A-E**) and MBP-immunoreactive myelin debris in the same part of damaged optic nerves in the glaucoma rat (**F-H**). Regarding the code “NR17L14” in A, “NR”, “17”, “L”, and “14” indicate “normal rat”, “case number 17”, “left optic nerve”, and “section number 14”, respectively. Concerning the code “GR3L10”, “GR” represents “glaucoma rat”. The particles and myelin debris were visualized by fluorescent immunohistochemistry using a mouse monoclonal anti-human myelin basic protein (MBPh) antibody (clone SMI-99; Covance, Princeton, NJ, USA; Alexa Fluor 594 label). These images were taken with an LSM700 confocal microscope (Carl Zeiss, Jena, Germany). Note that the bottom of each image is positioned at the border between the unmyelinated and myelinated regions. Additionally, each image of the optic nerve was split along the pia mater to remove the background from the measurement area. Scale bar = 250 µm in **H** for **A-G**. **I:** The image shows the orientation of **A-H**.

**FIGURE 8.**
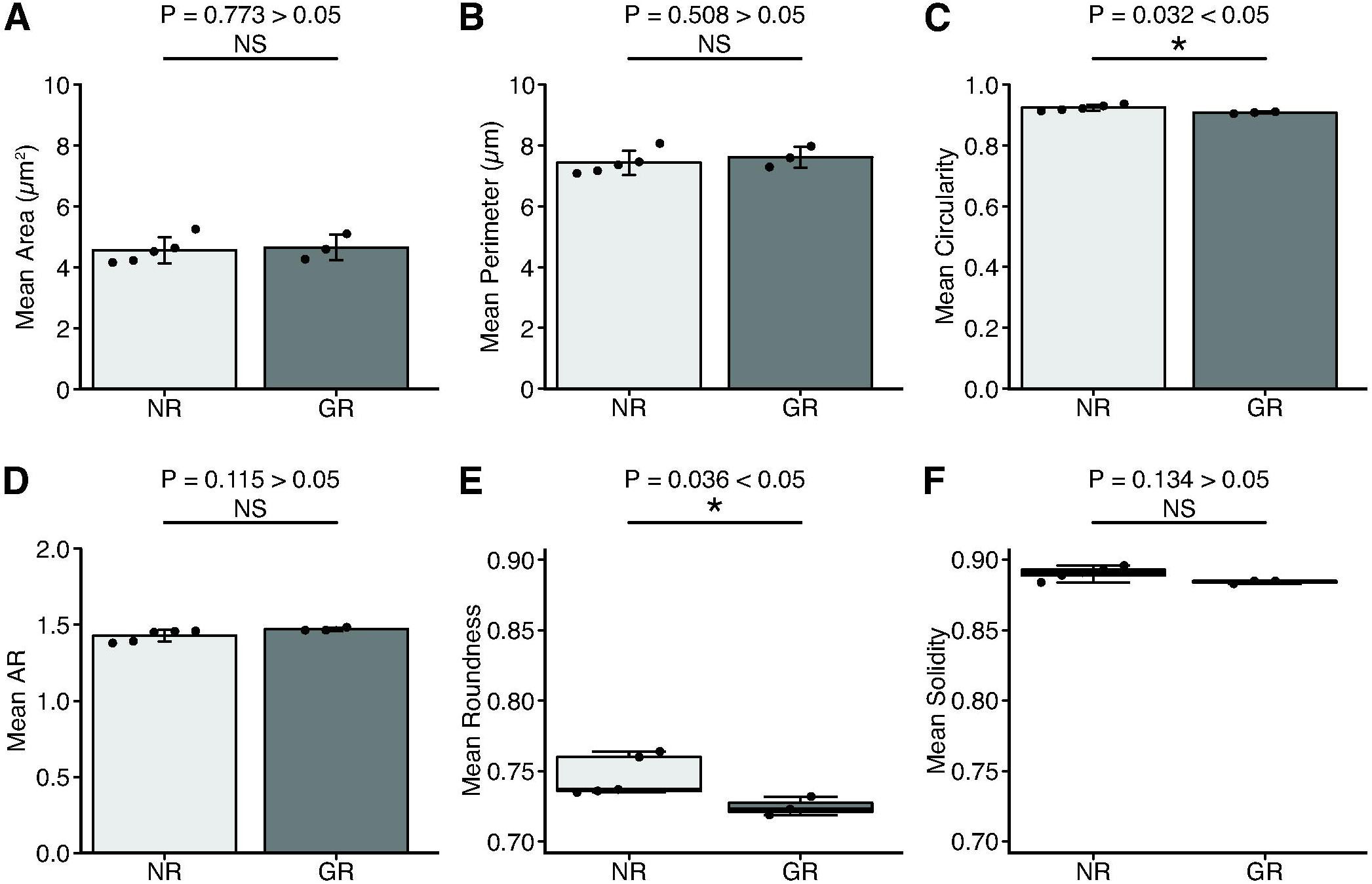
Charts show area, perimeter, and shape descriptors (circularity, AR (aspect ratio), roundness, and solidity) of myelin basic protein (MBP)-immunoreactive particles in the normal rat (NR) as well as those of myelin debris in damaged optic nerves of the glaucoma rat (GR). Images of the MBP-immunoreactive particles in the NR and those of the MBP-immunoreactive myelin debris in the GR were taken in the distal (anterior)-most part of the myelinated region of the optic nerve, as shown in Figure 7A-H. Since the anti-human myelin basic protein (MBPh) antibody (clone SMI-99) was used to visualize the particles and myelin debris, the term MBPh will be utilized instead of MBP in the following description. **A-D:** Data are presented as mean ± SD (standard deviation; n = 5 rats/ NR group; n = 3 rats/ GR group). Results of two-sample t-tests are indicated above the charts. An asterisk (*) in **C** represents a significant difference with P = 0.032 < 0.05. Each solid black circle in **A**, **B**, **C**, and in **D** denotes the mean area, mean perimeter, mean circularity, and mean AR of the MBPh-immunoreactive particles or those of MBPh-immunoreactive myelin debris in each experimental case, respectively. **A:** Mean area of MBPh-immunoreactive particles in the NR and that of MBPh-immunoreactive myelin debris in the GR; NR: mean ± SD = 4.562 ± 0.435; GR: mean ± SD = 4.657 ± 0.421. **B.** Mean perimeter; NR: mean ± SD = 7.425 ± 0.388; GR: mean ± SD = 7.617 ± 0.343. **C.** Mean circularity; NR: mean ± SD = 0.923 ± 0.009; GR: mean ± SD = 0.907 ± 0.003. **D.** Mean AR; NR: mean ± SD = 1.428 ± 0.039; GR: mean ± SD = 1.471 ± 0.011. **E-F:** Data are presented as box and whisker plots (n = 5 rats/ NR group; n = 3 rats/ GR group). The results of Mann-Whitney U tests are indicated above the charts. An asterisk (*) in **E** represents a significant difference with P = 0.036 < 0.05. Each solid black circle in **E**, and in **F** denotes the mean roundness and mean solidity of the MBPh-immunoreactive particles, or those of MBPh-immunoreactive myelin debris in each experimental case, respectively. The upper whisker indicates the maximum value, while the lower whisker represents the minimum value. The thick horizontal line in the box shows the median. The upper and lower borders of the box indicate the interquartile range. Note that the mean circularity and mean roundness of MBPh-immunoreactive particles were significantly different from those of MBPh-immunoreactive myelin debris (p < 0.05). Consequently, only 2 out of 6 differences in sizes and shape descriptors between the particles and the myelin debris were statistically significant. NS, not significant.

### 3.5 Distribution of MBPh-immunoreactive particles in the mouse and monkey optic nerves

A considerable number of MBPh-immunoreactive particles were distributed in the distal-most part of the myelinated region in the mouse optic nerve (Figure 9). In addition, a reasonable number of MBPh-immunoreactive particles were observed in the distal-most part of the retrolaminar region in the monkey optic nerve (Figure 10).

**FIGURE 9.**
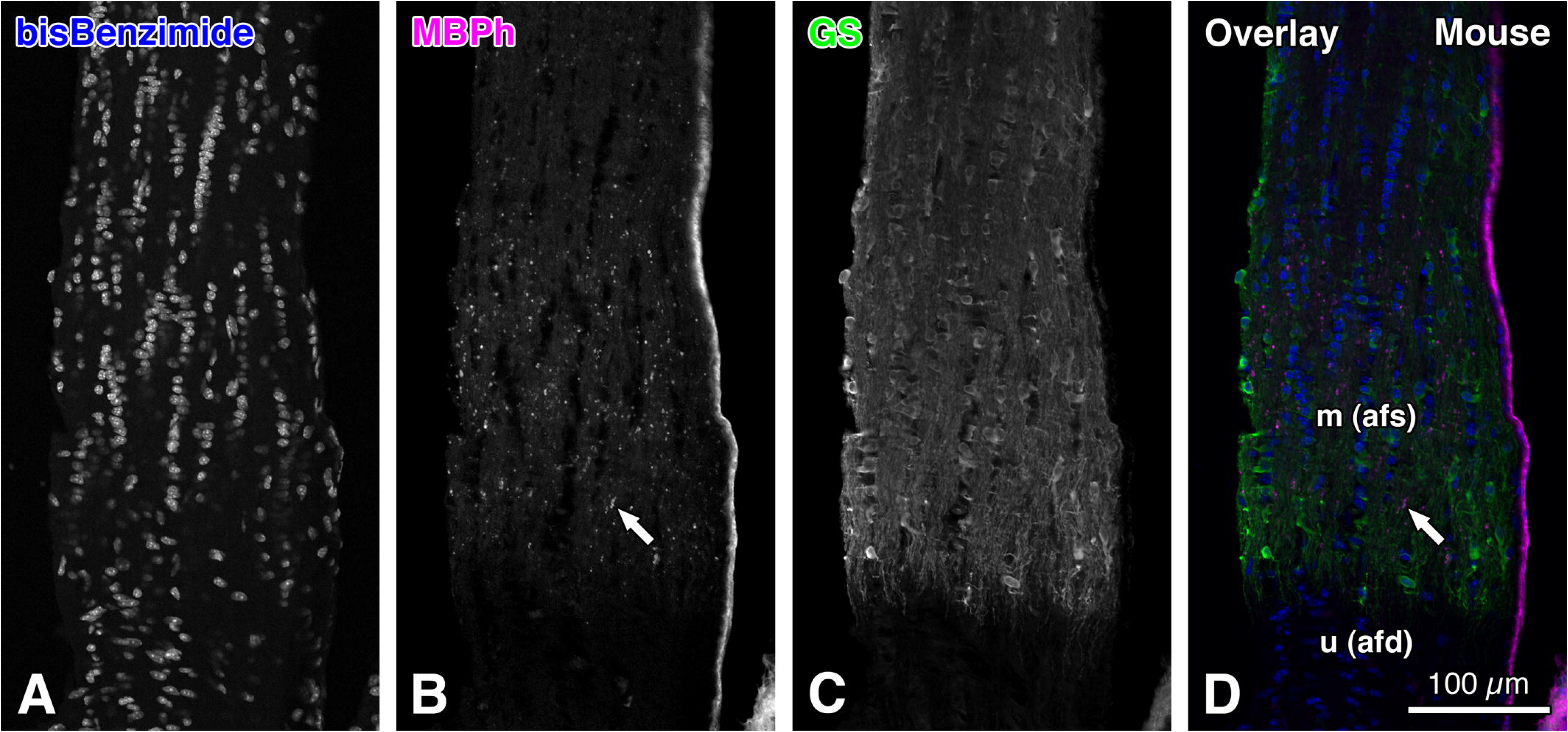
Images showing the distribution of myelin basic protein-immunoreactive particles in the myelinated region of the mouse optic nerve. The images represent the distribution of glutamine synthetase (GS) and of the particles visualized by using fluorescent double immunohistochemistry. The images were taken from a longitudinal section through the paramedian part in the distal (anterior)-most part of the myelinated region in the normal mouse optic nerve. The particles colored in white (**B**) and in magenta (**D**) were labeled with a mouse monoclonal anti-human myelin basic protein (MBPh) antibody (clone SMI-99; Covance, Princeton, NJ, USA; Alexa Fluor 594 label). The GS protein was labeled with an anti-GS antibody (Sigma-Aldrich, Saint Louis, MO, USA; Alexa Fluor 488 label; **C**; green in **D**). Cell nuclei were labeled with bisBenzimide (Hoechst 33258; **A**; blue in **D**). The arrows indicate MBPh-immunoreactive particles distributed in the myelinated region of the mouse optic nerve (**B**, **D**). These images were taken with an LSM 700 confocal microscope (Carl Zeiss, Jena, Germany). Note that a substantial number of MBPh-immunoreactive particles are distributed in the myelinated region of the mouse optic nerve. m (afs), myelinated region (astrocytic filament sparse region). u (afd), unmyelinated region (astrocytic filament dense region). Scale bar = 100 µm in the lower right corner of **D** for **A-C**.

**FIGURE 10.**
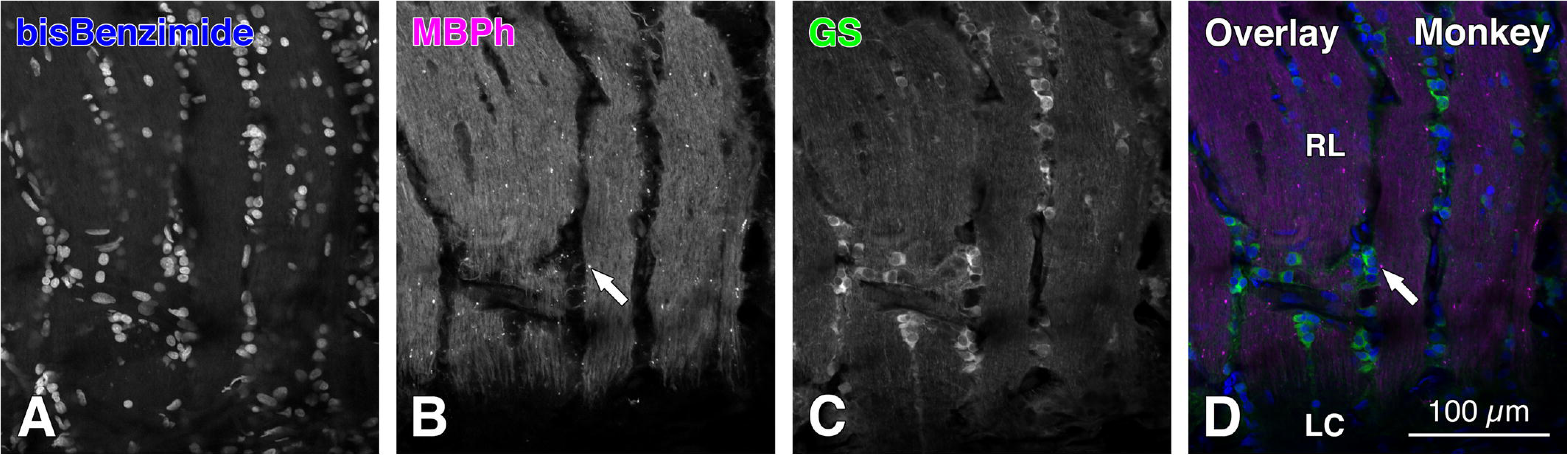
Images showing the distribution of myelin basic protein-immunoreactive particles in the retrolaminar (myelinated) region of the optic nerve in the monkey (*Macaca fuscata*). The images represent the distribution of glutamine synthetase (GS) and of the particles visualized by using fluorescent double immunohistochemistry. The images were taken from a longitudinal section through the paramedian part in the distal (anterior)-most part of the retrolaminar region in the monkey optic nerve. The particles colored in white (**B**) and in magenta (**D**) were labeled with a mouse monoclonal anti-human myelin basic protein (MBPh) antibody (clone SMI-99; Covance, Princeton, NJ, USA; Alexa Fluor 594 label). The GS protein was labeled with an anti-GS antibody (Sigma-Aldrich, Saint Louis, MO, USA; Alexa Fluor 488 label; **C**; green in **D**). Cell nuclei were labeled with bisBenzimide (Hoechst 33258; **A**; blue in **D**). In the monkey, myelinated fibers are clearly labeled with the anti-MBPh antibody. The arrows indicate an MBPh-immunoreactive particle distributed in the retrolaminar region of the monkey optic nerve (**B**, **D**). These images were taken with an LSM 700 confocal microscope (Carl Zeiss, Jena, Germany). Note that a substantial number of MBPh-immunoreactive particles are distributed in the retrolaminar region of the monkey optic nerve. LC, lamina cribrosa. RL, retrolaminar region. Scale bar = 100 µm in the lower right corner of **D** for **A-C**.

## 4 Discussion

### 4.1 A summary of the results

We demonstrated the concentration of MBPh-immunoreactive particles in the distal-most part of the myelinated region in the normal rat optic nerve. The MBPh-immunoreactive particles contained the authentic myelin basic protein. These particles colocalized with neural and glial cell marker proteins, such as GFAP, GS, and NF200 proteins. The majority of MBPh-immunoreactive particles were distributed on myelinated nerve fibers in the optic nerve and were isolated from Iba1-immunoreactive microglia. All differences in mean sizes (area and perimeter) and mean shape descriptors (circularity, AR (aspect ratio), roundness, and solidity) between the MBPh-immunoreactive particles and the MBPh-immunoreactive myelin debris in the damaged optic nerve of the glaucoma rat were less than 3 percent. The mean circularity and mean roundness of the particles were significantly different from those of the myelin debris. Accordingly, only 2 out of 6 differences in the mean sizes (area and perimeter) and the mean shape descriptors (circularity AR, roundness, and solidity) between the particles and the myelin debris were statistically significant. MBPh-immunoreactive particles were also observed in the distal-most part of the myelinated region in the mouse and monkey optic nerves.

### 4.2 The mouse anti-MBPh antibody (SMI-99)

#### 4.2.1 Specificity of the mouse anti-MBPh antibody (SMI-99)

The present study demonstrated the concentration of MBPh-immunoreactive particles in the distal-most part of the myelinated region in the normal rat optic nerve. Consequently, few reports have emerged regarding this evidence. Therefore, there exists a remote possibility that the particles were visualized due to a false-positive immunoreaction. It is necessary to examine whether the anti-MBPh antibody detected authentic MBP or a protein distinct from MBP. This notion was verified by using the anti-MBPc antibody; the target sequence of the anti-MBPc antibody differs from that of the anti-MBPh antibody (Saper 2005). Regarding the immunohistochemistry of the MBPh-immunoreactive particles, the anti-MBPh antibody against Ala-Ser-Asp-Tyr-Lys-Ser (ASDYKS) in position 131-136 of the classic human myelin basic protein (MBPh) demonstrated the same staining pattern as another anti-MBPc antibody against Asp-Glu-Asn-Pro-Val-Val (DENPVV) in position 82-87 of the full length protein of the cow myelin basic protein (MBPc; Figure 2). Furthermore, cross-reactions were not detectable between the mouse monoclonal anti-MBPh antibody and the Alexa Fluor 488 conjugated goat anti-rat secondary antibody (Spplementary Figure 1C), or between the rat monoclonal anti-MBPc antibody and the Alexa Fluor 594 conjugated goat anti-mouse secondary antibody (Supplementary Figure 1J). These facts indicate that the fluorescent double immunohistochemistry by using the mouse monoclonal anti-MBPh antibody and the rat monoclonal anti-MBPc antibody was not false-positive but genuine. Thus, the anti-MBPh antibody immunoreacted with the authentic MBP. Therefore, the MBPh-immunoreactive particles contained the authentic MBP.

#### 4.2.2 Detection of the MBPh-immunoreactive particles by using the mouse anti-MBPh antibody (SMI-99)

The MBP-immunoreactivity of the MBPh-immunoreactive particles, visualized by using the mouse anti-MBPh antibody (SMI-99) as the primary antibody, was equivalent to that visualized by using the rat anti-MBPc antibody (clone 12). However, the MBP-immunoreactivity of myelinated nerve fibers in the rat optic nerve, visualized by using the anti-MBPh antibody, was weaker than that visualized by using the anti-MBPc antibody (Figure 1). Consequently, the MBP-immunoreactivity of the MBPh-immunoreactive particles was much stronger than that of the MBPh-immunoreactive myelinated nerve fibers (Figure 1A). Therefore, we could clearly distinguish the particles from the myelinated nerve fibers by using the anti-MBPh antibody in the present study.

The MBP-immunoreactivity of the myelinated nerve fibers in the rat optic nerve, visualized by using the anti-MBPh antibody, was weaker than that in the monkey optic nerve (Figures 7A-E, 10B). Additionally, pig and chicken MBP do not react with the anti-MBPh antibody, and guinea pig MBP shows slight reactivity with this antibody (Manufacturer’s technical information). Therefore, the weak MBP-immunoreactivity of the rat myelinated nerve fibers, as visualized by using the anti-MBPh antibody, is attributable to species differences in the reactivity of the anti-MBPh antibody between the rat MBP protein and the monkey MBP protein.

Another underlying cause of the difference in MBPh-immunoreactivity between the particles and the myelinated nerve fibers can be attributed to a dissimilarity in the density of the antigen, MBP protein, present in each structure. The majority of the myelin debris-like MBPh-immunoreactive particles are considered to consist of broken myelin sheaths (Section 4.6). Consequently, these particles represent a structure rich in MBP. Conversely, the myelinated nerve fibers exhibit a lower concentration of MBP, since the cross-sectional area of the myelin sheath comprises approximately 40% of the cross-sectional area overall in the myelinated nerve fiber of the rat optic nerve (see footnote^3^). This evidence suggests that the density of MBP antigen in the MBPh-immunoreactive myelinated nerve fibers is less than half that found in the MBPh-immunoreactive particles. Therefore, the weaker MBPh-immunoreactivity in the myelinated nerve fibers can be attributable to the lower density of the antigen, the MBP protein, in the fibers.

### 4.3 Comparison with previous findings

MBP-immunoreactive myelin sheaths have been described in the myelinated region of the rodent optic nerve by several research groups (Dixon and Eng 1982; Morcos and Chan-Ling 2000; Sun et al. 2009). In the present study, we confirm these observations. Additionally, we demonstrate the concentration of MBPh-immunoreactive particles in the distal-most part of the myelinated region in the normal rat optic nerve. Few studies have reported this concentration. Therefore, the present study provides new evidence for myelin sheath morphology in the normal rat optic nerve.

### 4.4 Colocalization of neural and glial cell marker proteins in the MBPh-immunoreactive particles

In the distal-most part of the myelinated region, bundles of myelinated nerve fiber interdigitate with columns of GFAP-immunoreactive cells. Additionally, GFAP-immunoreactive filaments are aligned parallel to the optic nerve axis and are distributed among myelinated nerve fiber bundles (Morcos and Chan-Ling 2000; Kawano 2015b). In the present study, the MBPh-immunoreactive particles were observed in the distal-most part, and exhibited similarities to the MBPh-immunoreactive myelin debris found in the damaged optic nerve of the glaucoma rat (Section 4.6). It is reasonable to speculate that the GFAP-immunoreactivity in the MBPh-immunoreactive particles can be attributed to the GFAP-immunoreactive filaments located on or around myelinated nerve fibers, since the particles were composed of broken myelin sheaths (Section 4.6).

Myelin sheaths constitute components of oligodendrocytes (Peters et al. 1991; Bron et al. 1997), with 97 % of GS-immunoreactive cells identified as oligodendrocytes (Kawano 2015b). Additionally, GS-immunoreactive fibers extend longitudinally along the long axis of the rat optic nerve and are RIP-immunoreactive (Kawano 2015b). This evidence suggests that the myelin sheaths contain GS, since RIP is a marker of oligodendrocytes, including the myelin sheaths (Friedman et al. 1989). It is reasonable to infer that the GS-immunoreactivity observed in the MBPh-immunoreactive particles can be attributed to GS in the myelin sheaths, since these particles consist of broken myelin sheaths.

NF200-immunoreactive filaments were distributed in the axons of optic nerve neurons (Figure 4E-G; Peters et al. 1991). It is reasonable to speculate that NF200-immunoreactivity in the MBPh-immunoreactive particles is attributable to NF200 distributed in the axons. The particles likely involve parts of broken axons, since they were composed of broken myelin sheaths that had insulated the broken axons.

### 4.5 The majority of MBPh-immunoreactive particles were isolated from Iba1-immunoreactive microglia in the normal rat optic nerve

In the peripheral nervous system, the transection of a peripheral nerve leads to a prompt recruitment of hematogenous macrophages (Perry et al. 1987; Stoll et al. 1989a; Brück 1997; Stoll and Jander 1999). These macrophages migrate to degenerating nerve fibers and adhere to myelin ovoids containing myelin debris (Stoll et al. 1989a; Stoll and Jander 1999). Myelin debris is almost completely cleared within the first 2 weeks (Stoll and Jander 1999).

In the central nervous system, however, microglia are able to transform into large phagocytes, thereby removing myelin debris. Following the transection of the optic nerve or of fiber tracts in the spinal cord, there is an early transient period of microglial activation, but the appearance of large phagocytes is delayed by many weeks (Perry et al. 1987; Stoll et al. 1989b; George and Griffin 1994; Stoll and Jander 1999). Consequently, prolonged persistence of myelin debris in degenerating fiber tracts of the central nervous system has been observed (Stoll and Jander 1999; Weinger et al. 2011). This evidence suggests that the prolonged persistence of the MBPh-immunoreactive particles is possible in the normal rat optic nerve. Accordingly, the isolation of these particles from Iba1-immunoreactive microglia is also achievable, since there is little discrepancy between the isolation and the prolonged persistence of the MBP-immunoreactive particles.

### 4.6 Neuroanatomical comparison between the MBPh-immunoreactive particles and the MBPh-immunoreactive myelin debris

The MBPh-immunoreactive particles in the normal rat optic nerve were immunohistochemically similar to the MBPh-immunoreactive myelin debris in the damaged optic nerve of the glaucoma rat. This notion is supported by four facts. Firstly, both the MBPh-immunoreactive particles and myelin debris exhibited immunohistochemistry for myelin basic protein (MBP). Secondly, the size of the MBPh-immunoreactive particles was broadly similar to that of the MBPh-immunoreactive myelin debris (Figures 6-7). Thirdly, all differences in the mean sizes (area and perimeter) and the mean shape descriptors (circularity, AR (aspect ratio), roundness, and solidity) between the particles and the myelin debris were less than 3 percent (Table 2). Fourthly, the mean circularity and mean roundness of the particles were significantly different from those of the myelin debris. Accordingly, only 2 out of 6 differences in the mean sizes and the mean shape descriptors between the particles and the myelin debris were statistically significant (Figure 8). Thus, subtle but significant differences between the particles and the myelin debris were detected in two mean shape descriptors: mean circularity and mean roundness. Based on these facts, it is extremely difficult to categorize the particles and myelin debris separately. Therefore, it is possible to accept that the MBPh-immunoreactive particles were immunohistochemically similar to the MBPh-immunoreactive myelin debris. Probably, the particles consisted of broken myelin sheaths. This notion is supported by evidence that the majority of MBPh-immunoreactive particles were distributed on myelinated nerve fibers in both the normal and glaucoma rat optic nerves (Figures 1A, 6A, 7A-H). Accordingly, it is appropriate to describe the particles as myelin debris-like MBPh-immunoreactive particles.

### 4.7 Distribution of MBPh-immunoreactive particles in the optic nerves of the mouse, rat, and monkey

The distribution of MBPh-immunoreactive particles was observed not only in the rat but also in the mouse and monkey optic nerves in their distal-most parts (Figures 1A, 9, 10). These findings suggest that the distribution of MBPh-immunoreactive particles in the optic nerve is similar among various mammalian species.

### 4.8 Underlying causes of the concentration of MBPh-immunoreactive particles in the distal-most part of the myelinated region in the normal rat optic nerve

The underlying causes of the concentration are discussed in detail in the companion paper by Kawano (2026). In brief, recent evidence indicates that disruption of energy in neurons and glia leads to myelin damage (Micu et al. 2006; Hamilton et al. 2016; Huang et al. 2023; Anderle et al. 2025). During severe energy deprivation, the myelin sheath sustains damage due to the activation of Ca^2+^ fluxes through NMDA (*N*-methyl-D-aspartate) receptors (Micu et al. 2006) and TRPA1 (transient receptor potential ankyrin 1) channels (Hamilton et al. 2016; Anderle et al. 2025). Accordingly, there is a possibility that the concentration of the MBPh-immunoreactive particles in the distal-most part is attributable to the lower level of energy supply to this region due to the smaller amount of microvascular blood flow in this region (Kawano 2026).

GFAP and GS are abundantly distributed in the distal-most part (Kawano 2026). Since reactive astrogliosis is a fibrous proliferation of glial cells in injured areas of the central nervous system, the abundant distribution of GFAP in the distal-most part suggests that this area might be under physiologically stressed conditions (Messam et al. 2002; McAteer and Choudhury 2009; Sofroniew 2009; Kawano 2026). Given that GS in oligodendrocytes increases in chronic pathological conditions in both mice and humans (Ben Haim et al. 2021), the abundant distribution of GS in the distal-most part suggests that this area might be under physiologically stressed conditions (Kawano 2026). It is possible that these histopathological backgrounds in the distal-most part impair myelin sheaths, subsequently leading to the formation of myelin debris-like MBPh-immunoreactive particles.

### 4.9 The contribution of the MBPh-immunoreactive particles to neurohistopathology in the future

The concentration of MBPh-immunoreactive particles was observed in the distal-most part of the myelinated region in the normal rat optic nerve. These particles were immunohistochemically similar to MBPh-immunoreactive myelin debris found in the damaged optic nerve of the glaucoma rat (Section 4.6). Therefore, the damaged optic nerve was in a severe pathological condition. The density of MBPh-immunoreactive myelin debris in the damaged optic nerve of the glaucoma rat was higher than that in the distal-most part of the normal rat (Figure 7). Additionally, the density of these particles in the distal-most part is the highest in the myelinated region of the normal rat (Kawano 2026). Recently, the distal-most part of the myelinated region has been considered to be in a physiologically stressed condition, while the other parts remain in physiological conditions (Kawano 2026). Thus, the density of MBPh-immunoreactive particles and/or myelin debris varied according to histopathological conditions. Therefore, it is possible that MBPh-immunoreactive particles and/or myelin debris can serve as histopathological biomarkers.

## 5 Conclusion

In summary, we demonstrated the concentration of MBPh-immunoreactive particles in the distal-most part of the myelinated region in the normal rat optic nerve. The MBPh-immunoreactive particles contained authentic myelin basic protein. These particles were morphologically similar to the MBPh-immunoreactive myelin debris found in the damaged optic nerve of the glaucoma rat. Additionally, MBPh-immunoreactive particles were also observed in the distal-most part of the myelinated region in the optic nerves of both the mouse and monkey.

These results demonstrate that the myelin debris-like MBPh-immunoreactive particles are concentrated in the distal-most part of the myelinated region. This evidence suggests that this area is in a physiologically stressed condition. Additionally, this evidence may provide valuable insights into the pathophysiological mechanisms that induce localized vulnerability of the myelin sheaths. Furthermore, it is possible that MBPh-immunoreactive particles and/or myelin debris can serve as histopathological biomarkers, since the density of these particles and/or myelin debris varies according to histopathological conditions (Section 4.9).

## Author Contributions

J.K.: Concept, Funding, Supervision, Investigation, Data analysis, Writing original draft, Writing – review and editing.

## Acknowledgments

The author thanks Doctor Shiro Nakagawa (Professor Emeritus, Kagoshima University Graduate School of Medical and Dental Sciences) for supplying the monkey eyes and optic nerves, Professor Masahisa Horiuchi (Department of Hygiene and Health Promotion Medicine, Kagoshima University Graduate School of Medical and Dental Sciences) for his expert guidance on statistical analyses, and Associate Professor Kentaro Setoyama (Division of Laboratory Animal Resources and Research, Center for Advanced Science Research and Promotion, Kagoshima University) for his expert advice on managing glaucoma rats. This work was supported by an annual fund from Kagoshima University.

## Data Availability Statement

The data that support the findings of this study are available from the corresponding author upon reasonable request.

## Patient Consent Statement

Not applicable.

## Permission to Reproduce Material from Other Sources

Not applicable.

## Clinical Trial Registration

Not applicable.

---

**SUPPLEMENTARY FIGURE 1.**
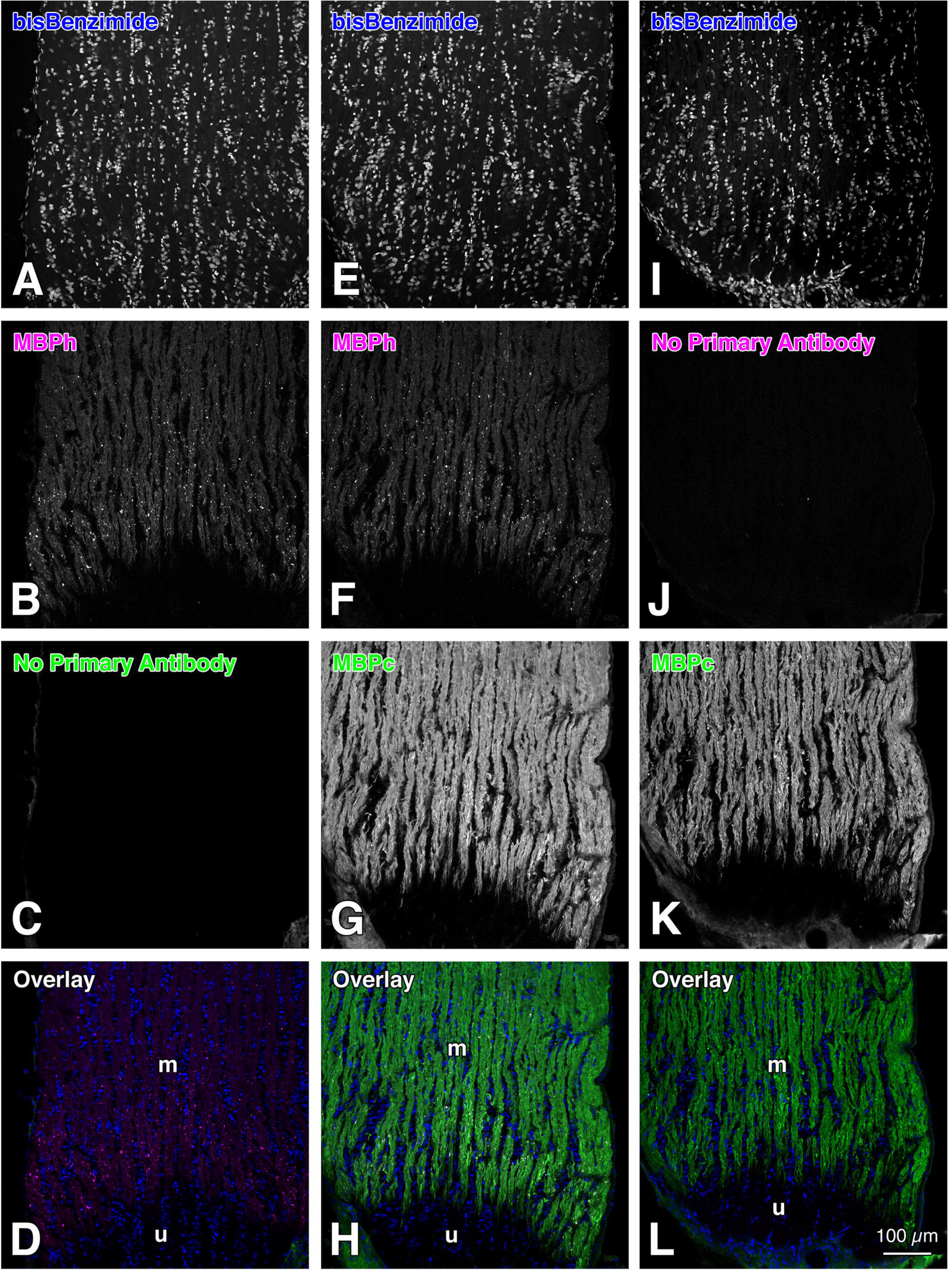
Images of fluorescent double immunohistochemistry on three serial optic nerve sections demonstrating undetectable cross-reactions between a mouse monoclonal primary antibody and a goat anti-rat secondary antibody (**C**), as well as between a rat monoclonal primary antibody and a goat anti-mouse secondary antibody (**J**). The two monoclonal primary antibodies used were anti-myelin basic protein (MBP) antibodies targeting different amino acid sequences. The three serial optic nerve sections are longitudinal sections through the paramedian part in the distal (anterior)-most part of the myelinated region in the normal rat optic nerve. Additionally, panels **A-D** show the first section, panels **E-H** indicate the second section, and panels **I-L** demonstrate the third section of the three serial optic nerve sections. **A, E, I:** Cell nuclei labeled with bisBenzimide (Hoechst 33258; blue in **D**, **H**, and **L**). **B, F:** Particles and myelinated nerve fibers were labeled with the mouse monoclonal anti-human myelin basic protein (MBPh) antibody (clone SMI-99; Covance, Princeton, NJ, USA; Alexa Fluor 594 label; magenta in **D**, and in **H**). This antibody reacts with Ala-Ser-Asp-Tyr-Lys-Ser (ASDYKS) in position 131-136 of the classic human myelin basic protein. **C:** No primary antibody generated in the rat (the rat monoclonal anti-MBPc antibody, see **G**, **K**) was applied. **D, H, L:** Panels **D**, **H**, and **L** are overlay images of panels **A-C**, **E-G**, and **I-K**, respectively. **J:** No primary antibody generated in the mouse (the mouse monoclonal anti-MBPh antibody, see **B**, **F**) was applied. **G, K:** Particles and myelinated nerve fibers were labeled with the rat monoclonal anti-cow myelin basic protein (MBPc) antibody (clone 12; Abcam, Cambridge, United Kingdom; Alexa Fluor 488 label; green in **H**, and in **L**). This antibody reacts with Asp-Glu-Asn-Pro-Val-Val (DENPVV) in position 82-87 of the full length protein of cow myelin basic protein. These images were taken with an LSM 700 confocal microscope (Carl Zeiss, Jena, Germany). Note that cross-reactions are not detectable between the mouse monoclonal anti-MBPh antibody and the Alexa Fluor 488 conjugated goat anti-rat secondary antibody (**C**), or between the rat monoclonal anti-MBPc antibody and the Alexa Fluor 594 conjugated goat anti-mouse secondary antibody (**J**). These facts indicate that the fluorescent double immunohistochemistry by using the mouse monoclonal anti-MBPh antibody and the rat monoclonal anti-MBPc antibody was authentic rather than artificial. Consequently, the MBPh and MBPc double-immunoreactive particles visualized in Figure 2D were authentic rather than false positives. m, myelinated region; u, unmyelinated region. Scale bar = 100 µm in the lower right corner of **L** for **A-K**.

## Footnotes

^1^The GS/MBPh staining indicates double-label immunohistochemistry by using the rabbit polyclonal anti-GS antibody and the mouse monoclonal anti-MBPh antibody.

^2^The MBPc control staining shows double-label immunohistochemistry by using the 10% normal goat serum (NGS) blocking solution proper instead of the mouse monoclonal anti-MBPh antibody solution in the first step, followed by using the rat monoclonal anti-MBPc antibody solution in the next step.

^3^Forrester and Peters (1967) measured the diameters of myelinated nerve fibers in the optic nerve of the albino rat. The modal diameter overall is 0.9 µm. The modal diameter, excluding the myelin sheath, is 0.7 µm. Based on this evidence, the author calculated the following values of a myelinated nerve fiber: the cross-sectional area of the myelin sheath was 0.2512 µm^2^ (39.51%); the cross-sectional area excluding the myelin sheath was (0.35 X 0.35 X 3.14=) 0.38465 µm^2^ (60.49%); and the cross-sectional area overall was (0.45 X 0.45 X 3.14 =) 0.63585 µm^2^ (100.00%).

## References

1. Anderle, S., Dixon, M., Quintela-Lopez, T., Sideris-Lampretsas, G., & Attwell, D. (2025). The vascular contribution to cognitive decline in ageing and dementia. Nat Rev Neurosci, 26(10), 591–606. 10.1038/s41583-025-00950-1

2. Balaratnasingam, C., Morgan, W. H., Johnstone, V., Cringle, S. J., & Yu, D. Y. (2009). Heterogeneous distribution of axonal cytoskeleton proteins in the human optic nerve. Investigative ophthalmology & visual science, 50(6), 2824–2838. 10.1167/iovs.08-3206

3. Ben Haim, L., Schirmer, L., Zulji, A., Sabeur, K., Tiret, B., Ribon, M., Chang, S., Lamers, W. H., Boillée, S., Chaumeil, M. M., & Rowitch, D. H. (2021). Evidence for glutamine synthetase function in mouse spinal cord oligodendrocytes. Glia, 69(12), 2812–2827. 10.1002/glia.24071

4. Benvegnù, S., Poggiolini, I., & Legname, G. (2010). Neurodevelopmental expression and localization of the cellular prion protein in the central nervous system of the mouse. The Journal of Comparative Neurology, 518(11), 1879–1891. 10.1002/cne.22357

5. Black, J. A., Waxman, S. G., & Hildebrand, C. (1985). Axo-glial relations in the retina-optic nerve junction of the adult rat: freeze-fracture observations on axon membrane structure. Journal of Neurocytology, 14(6), 887–907. 10.1007/BF01224803

6. Bosco, A., Inman, D. M., Steele, M. R., Wu, G., Soto, I., Marsh-Armstrong, N., Hubbard, W. C., Calkins, D. J., Horner, P. J., & Vetter, M. L. (2008). Reduced retina microglial activation and improved optic nerve integrity with minocycline treatment in the DBA/2J mouse model of glaucoma. Investigative ophthalmology & visual science, 49(4), 1437–1446. 10.1167/iovs.07-1337

7. Bosco, A., Steele, M. R., & Vetter, M. L. (2011). Early microglia activation in a mouse model of chronic glaucoma. The Journal of Comparative Neurology, 519(4), 599–620. 10.1002/cne.22516

8. Bron, A. J., Tripathi, R. C., & Tripathi, B. J. (1997). The Visual Pathway. In: A. J. Bron, R. C. Tripathi, & B. J. Tripathi (Eds.), Wolff’s Anatomy of the Eye and Orbit. (8th ed., pp. 489–596). London: Chapman & Hall.

9. Brück, W. (1997). The role of macrophages in Wallerian degeneration. Brain Pathology, 7(2), 741–752. 10.1111/j.1750-3639.1997.tb01060.x

10. Butt, A. M. (2013). Structure and function of oligodendrocytes. In H. Kettenmann & B. R. Ransom (Eds.), Neuroglia (3rd ed., pp. 62–73). New York: Oxford University Press.

11. Butt, A. M., & Ransom, B. R. (1993). Morphology of astrocytes and oligodendrocytes during development in the intact rat optic nerve. The Journal of Comparative Neurology, 338(1), 141–158. 10.1002/cne.903380110

12. Chang, M. L., Wu, C. H., Jiang-Shieh, Y. F., Shieh, J. Y., & Wen, C. Y. (2007). Reactive changes of retinal astrocytes and Müller glial cells in kainate-induced neuroexcitotoxicity. Journal of Anatomy, 210(1), 54–65. 10.1111/j.1469-7580.2006.00671.x

13. Chen, M. S., Huber, A. B., van der Haar, M. E., Frank, M., Schnell, L., Spillmann, A. A., Christ, F., & Schwab, M. E. (2000). Nogo-A is a myelin-associated neurite outgrowth inhibitor and an antigen for monoclonal antibody IN-1. Nature, 403(6768), 434–439. 10.1038/35000219

14. Chrast, R., Saher, G., Nave, K. A., & Verheijen, M. H. (2011). Lipid metabolism in myelinating glial cells: lessons from human inherited disorders and mouse models. Journal of Lipid Research, 52(3), 419–434. 10.1194/jlr.R009761

15. Dixon, R. G., & Eng, L. F. (1982). Processing techniques for the demonstration of myelin basic protein in paraffin-embedded optic nerve: an immunoperoxidase study of the developing albino rat. Journal of Histochemistry and Cytochemistry, 30(3), 270–273. 10.1177/30.3.6174565

16. Dugas, J. C., Tai, Y. C., Speed, T. P., Ngai, J., & Barres, B. A. (2006). Functional genomic analysis of oligodendrocyte differentiation. Journal of Neuroscience, 26(43), 10967–10983. 10.1523/jneurosci.2572-06.2006

17. Duncan, I. D., & Radcliff, A. B. (2016). Inherited and acquired disorders of myelin: The underlying myelin pathology. Experimental Neurology, 283(Pt B), 452–475. 10.1016/j.expneurol.2016.04.002

18. Dyer, C. A., Kendler, A., Philibotte, T., Gardiner, P., Cruz, J., & Levy, H. L. (1996). Evidence for central nervous system glial cell plasticity in phenylketonuria. Journal of Neuropathology & Experimental Neurology, 55(7), 795–814. 10.1097/00005072-199607000-00005

19. Filbin, M. T. (2003). Myelin-associated inhibitors of axonal regeneration in the adult mammalian CNS. Nat Rev Neurosci, 4(9), 703–713. 10.1038/nrn1195

20. Forrester, J., & Peters, A. (1967). Nerve fibres in optic nerve of rat. Nature, 214(5085), 245–247. 10.1038/214245a0

21. Fox, J. (2017). Using the R commander, A Point-and-Click Interface for R (1st ed.). Boca Raton, Florida: Chapman and Hall/CRC Press. 10.1201/9781315380537

22. Friedman, B., Hockfield, S., Black, J. A., Woodruff, K. A., & Waxman, S. G. (1989). In situ demonstration of mature oligodendrocytes and their processes: An immunocytochemical study with a new monoclonal antibody, Rip. Glia, 2(5), 380–390. 10.1002/glia.440020510

23. Gaillard, F., Bonfield, S., Gilmour, G. S., Kuny, S., Mema, S. C., Martin, B. T., Smale, L., Crowder, N., Stell, W. K., & Sauvé, Y. (2008). Retinal anatomy and visual performance in a diurnal cone-rich laboratory rodent, the Nile grass rat (*Arvicanthis niloticus*). The Journal of Comparative Neurology, 510(5), 525–538. 10.1002/cne.21798

24. George, R., & Griffin, J. W. (1994). Delayed macrophage responses and myelin clearance during Wallerian degeneration in the central nervous system: the dorsal radiculotomy model. Experimental Neurology, 129(2), 225–236. 10.1006/exnr.1994.1164

25. Hamilton, N. B., Kolodziejczyk, K., Kougioumtzidou, E., & Attwell, D. (2016). Proton-gated Ca^2+^-permeable TRP channels damage myelin in conditions mimicking ischaemia. Nature, 529(7587), 523–527. 10.1038/nature16519

26. Haverkamp, S., & Wässle, H. (2000). Immunocytochemical analysis of the mouse retina. The Journal of Comparative Neurology, 424(1), 1–23. 10.1002/1096-9861(20000814)424:1<1::AID-CNE1>3.0.CO;2-V

27. Hojo, M., Ohtsuka, T., Hashimoto, N., Gradwohl, G., Guillemot, F., & Kageyama, R. (2000). Glial cell fate specification modulated by the bHLH gene *Hes5* in mouse retina. Development, 127(12), 2515–2522. 10.1242/dev.127.12.2515

28. Howell, G. R., Libby, R. T., Jakobs, T. C., Smith, R. S., Phalan, F. C., Barter, J. W., Barbay, J. M., Marchant, J. K., Mahesh, N., Porciatti, V., Whitmore, A. V., Masland, R. H., & John, S. W. (2007). Axons of retinal ganglion cells are insulted in the optic nerve early in DBA/2J glaucoma. Journal of Cell Biology, 179(7), 1523–1537. 10.1083/jcb.200706181

29. Huang, S., Ren, C., Luo, Y., Ding, Y., Ji, X., & Li, S. (2023). New insights into the roles of oligodendrocytes regulation in ischemic stroke recovery. Neurobiol Dis, 184, 106200. 10.1016/j.nbd.2023.106200

30. Imai, Y., Ibata, I., Ito, D., Ohsawa, K., & Kohsaka, S. (1996). A novel gene iba1 in the major histocompatibility complex class III region encoding an EF hand protein expressed in a monocytic lineage. Biochemical and Biophysical Research Communications, 224(3), 855–862. 10.1006/bbrc.1996.1112

31. Ito, D., Tanaka, K., Suzuki, S., Dembo, T., & Fukuuchi, Y. (2001). Enhanced expression of Iba1, ionized calcium-binding adapter molecule 1, after transient focal cerebral ischemia in rat brain. Stroke, 32(5), 1208–1215. 10.1161/01.str.32.5.1208

32. Jeon, S. B., Yoon, H. J., Park, S. H., Kim, I. H., & Park, E. J. (2008). Sulfatide, a major lipid component of myelin sheath, activates inflammatory responses as an endogenous stimulator in brain-resident immune cells. Journal of Immunology, 181(11), 8077–8087. 10.4049/jimmunol.181.11.8077

33. Johnson, E. C., Jia, L., Cepurna, W. O., Doser, T. A., & Morrison, J. C. (2007). Global changes in optic nerve head gene expression after exposure to elevated intraocular pressure in a rat glaucoma model. Investigative ophthalmology & visual science, 48(7), 3161–3177. 10.1167/iovs.06-1282

34. Ju, W. K., Misaka, T., Kushnareva, Y., Nakagomi, S., Agarwal, N., Kubo, Y., Lipton, S. A., & Bossy-Wetzel, E. (2005). OPA1 expression in the normal rat retina and optic nerve. The Journal of Comparative Neurology, 488(1), 1–10. 10.1002/cne.20586

35. Kanamori, A., Nakamura, M., Nakanishi, Y., Nagai, A., Mukuno, H., Yamada, Y., & Negi, A. (2004). Akt is activated via insulin/IGF-1 receptor in rat retina with episcleral vein cauterization. Brain Research, 1022(1-2), 195–204. 10.1016/j.brainres.2004.06.077

36. Kanda, Y. (2013). Investigation of the freely available easy-to-use software ’EZR’ for medical statistics. Bone Marrow Transplantation, 48(3), 452–458. 10.1038/bmt.2012.244

37. Kawano, J. (2015a). Chemoarchitecture of glial fibrillary acidic protein (GFAP) and glutamine synthetase in the optic nerve of the monkey (*Macaca fuscata*): An immunohistochemical study. Okajimas Folia Anatomica Japonica, 91(4), 97–104. 10.2535/ofaj.91.97

38. Kawano, J. (2015b). Chemoarchitecture of glial fibrillary acidic protein (GFAP) and glutamine synthetase in the rat optic nerve: An immunohistochemical study. Okajimas Folia Anatomica Japonica, 92(1), 11–30. 10.2535/ofaj.92.11

39. Kawano, J. (2026). The heterogeneous distribution of glial cell marker proteins in the myelinated region of the normal rat optic nerve. bioRxiv, 10.1101/2025.03.19.643607

40. Kawano, J., Tanizawa, Y., & Shinoda, K. (2008). Wolfram syndrome 1 (*Wfs1*) gene expression in the normal mouse visual system. The Journal of Comparative Neurology, 510(1), 1–23. 10.1002/cne.21734

41. Kotter, M. R., Li, W. W., Zhao, C., & Franklin, R. J. (2006). Myelin impairs CNS remyelination by inhibiting oligodendrocyte precursor cell differentiation. Journal of Neuroscience, 26(1), 328–332. 10.1523/jneurosci.2615-05.2006

42. McAteer, M. A., & Choudhury, R. P. (2009). Applications of nanotechnology in molecular imaging of the brain. Progress in Brain Research, 180, 72–96. 10.1016/s0079-6123(08)80004-0

43. Melo, P., Moreno, V. Z., Vázquez, S. P., Pinazo-Durán, M. D., & Tavares, M. A. (2006). Myelination changes in the rat optic nerve after prenatal exposure to methamphetamine. Brain Research, 1106(1), 21–29. 10.1016/j.brainres.2006.05.020

44. Messam, C., Hou, J., Janabi, N., Monaco, M., Gravell, M., & Major, E. (2002). Glial cell types. In V. S. Ramachandran (Ed.), Encyclopedia of the Human Brain (1st ed., Vol. 2, pp. 369–387). San Diego: Academic Press. 10.1016/B0-12-227210-2/00152-7

45. Micu, I., Jiang, Q., Coderre, E., Ridsdale, A., Zhang, L., Woulfe, J., Yin, X., Trapp, B.D., McRory, J.E., Rehak, R., Zamponi, G.W., Wang, W., & Stys, P.K. (2006). NMDA receptors mediate calcium accumulation in myelin during chemical ischaemia. Nature, 439(7079), 988–992. 10.1038/nature04474

46. Morcos, Y., & Chan-Ling, T. (2000). Concentration of astrocytic filaments at the retinal optic nerve junction is coincident with the absence of intra-retinal myelination: comparative and developmental evidence. Journal of Neurocytology, 29(9), 665–678. 10.1023/A:1010835404754

47. Naka, M., Kanamori, A., Negi, A., & Nakamura, M. (2010). Reduced expression of aquaporin-9 in rat optic nerve head and retina following elevated intraocular pressure. Investigative ophthalmology & visual science, 51(9), 4618–4626. 10.1167/iovs.09-4712

48. Naskar, R., Wissing, M., & Thanos, S. (2002). Detection of early neuron degeneration and accompanying microglial responses in the retina of a rat model of glaucoma. Investigative ophthalmology & visual science, 43(9), 2962–2968.

49. Perry, V. H., Brown, M. C., & Gordon, S. (1987). The macrophage response to central and peripheral nerve injury. A possible role for macrophages in regeneration. Journal of Experimental Medicine, 165(4), 1218–1223. 10.1084/jem.165.4.1218

50. Peters, A., Palay, S. L., & Webster, H. d. (1991). The Fine Structure of the Nervous System: Neurons and Their Supporting Cells (3rd ed.). New York: Oxford University Press.

51. Riepe, R. E., & Norenburg, M. D. (1977). Müller cell localisation of glutamine synthetase in rat retina. Nature, 268(5621), 654–655. 10.1038/268654a0

52. Saari, J. C., Huang, J., Possin, D. E., Fariss, R. N., Leonard, J., Garwin, G. G., Crabb, J. W., & Milam, A. H. (1997). Cellular retinaldehyde-binding protein is expressed by oligodendrocytes in optic nerve and brain. Glia, 21(3), 259–268. 10.1002/(SICI)1098-1136(199711)21:3<259::AID-GLIA1>3.0.CO;2-0

53. Santos, A. M., Calvente, R., Tassi, M., Carrasco, M. C., Martín-Oliva, D., Martín -Teva, J. L., Navascués J., & Cuadros, M. A. (2008). Embryonic and postnatal development of microglial cells in the mouse retina. The Journal of Comparative Neurology, 506(2), 224–239. 10.1002/cne.21538

54. Saper, C. B. (2005). An open letter to our readers on the use of antibodies. The Journal of Comparative Neurology, 493(4), 477–478. 10.1002/cne.20839

55. Shareef, S. R., Garcia-Valenzuela, E., Salierno, A., Walsh, J., & Sharma, S. C. (1995). Chronic ocular hypertension following episcleral venous occlusion in rats. Experimental Eye Research, 61(3), 379–382. 10.1016/s0014-4835(05)80131-9

56. Smith, S. B., Brodjian, S., Desai, S., & Sarthy, V. (1997). Glial fibrillary acidic protein (GFAP) is synthesized in the early stages of the photoreceptor cell degeneration of the *mi*^vit^*/mi*^vit^ (vitiligo) mouse. Experimental Eye Research, 64(4), 645–650. 10.1006/exer.1996.0249

57. Sofroniew, M. V. (2009). Molecular dissection of reactive astrogliosis and glial scar formation. Trends Neurosci, 32(12), 638–647. doi:10.1016/j.tins.2009.08.002

58. Stoll, G., & Jander, S. (1999). The role of microglia and macrophages in the pathophysiology of the CNS. Progress in Neurobiology, 58(3), 233–247. 10.1016/s0301-0082(98)00083-5

59. Stoll, G., Griffin, J. W., Li, C. Y., & Trapp, B. D. (1989a). Wallerian degeneration in the peripheral nervous system: participation of both Schwann cells and macrophages in myelin degradation. Journal of Neurocytology, 18(5), 671–683. 10.1007/bf01187086

60. Stoll, G., Trapp, B. D., & Griffin, J. W. (1989b). Macrophage function during Wallerian degeneration of rat optic nerve: clearance of degenerating myelin and Ia expression. Journal of Neuroscience, 9(7), 2327–2335. 10.1523/jneurosci.09-07-02327.1989

61. Sun, D., Lye-Barthel, M., Masland, R. H., & Jakobs, T. C. (2009). The Morphology and Spatial Arrangement of Astrocytes in the Optic Nerve Head of the Mouse. The Journal of Comparative Neurology, 516(1), 1–19. 10.1002/cne.22058

62. Sun, X., Wang, X., Chen, T., Li, T., Cao, K., Lu, A., Chen, Y., Sun, D., Luo, J., Fan, J., Young, W., & Ren, Y. (2010). Myelin activates FAK/Akt/NF-kappaB pathways and provokes CR3-dependent inflammatory response in murine system. PLoS One, 5(2), e9380. 10.1371/journal.pone.0009380

63. Syed, Y. A., Zhao, C., Mahad, D., Möbius, W., Altmann, F., Foss, F., González, G. A., Sentürk, A., Acker-Palmer, A., Lubec, G., Lilley, K., Franklin, R. J. M., Nave, K. A., & Kotter, M. R. N. (2016). Antibody-mediated neutralization of myelin-associated EphrinB3 accelerates CNS remyelination. Acta Neuropathologica, 131(2), 281–298. doi:10.1007/s00401-015-1521-1

64. Talos, D. M., Fishman, R. E., Park, H., Folkerth, R. D., Follett, P. L., Volpe, J. J., & Jensen, F. E. (2006). Developmental regulation of alpha-amino-3-hydroxy-5-methyl-4-isoxazole-propionic acid receptor subunit expression in forebrain and relationship to regional susceptibility to hypoxic/ischemic injury. I. Rodent cerebral white matter and cortex. The Journal of Comparative Neurology, 497(1), 42–60. 10.1002/cne.20972

65. Wang, X., Cao, K., Sun, X., Chen, Y., Duan, Z., Sun, L., Guo, L., Bai, P., Sun, D., Fan, J., He, X., Young, W., & Ren, Y. (2015). Macrophages in spinal cord injury: phenotypic and functional change from exposure to myelin debris. Glia, 63(4), 635–651. 10.1002/glia.22774

66. Warr, W. B., de Olmos, J. S., & Heimer, L. (1981). Horseradish Peroxidase: The Basic Procedure. In L. Heimer & M. J. Robards (Eds.), Neuroanatomical tract-tracing methods. (1st ed., pp. 207–262). New York: Plenum Press.

67. Weinger, J. G., Brosnan, C. F., Loudig, O., Goldberg, M. F., Macian, F., Arnett, H. A., Prieto, A. L., Tsiperson, V., & Shafit-Zagardo, B. (2011). Loss of the receptor tyrosine kinase Axl leads to enhanced inflammation in the CNS and delayed removal of myelin debris during Experimental Autoimmune Encephalomyelitis. Journal of Neuroinflammation, 8, 49. 10.1186/1742-2094-8-49

68. Ye, H., & Hernandez, M. R. (1995). Heterogeneity of astrocytes in human optic nerve head. The Journal of Comparative Neurology, 362(4), 441–452. 10.1002/cne.903620402

69. Zabouri, N., Bouchard, J.-F., & Casanova, C. (2011). Cannabinoid receptor type 1 expression during postnatal development of the rat retina. The Journal of Comparative Neurology, 519(7), 1258–1280. 10.1002/cne.22534

70. Zhang, C., Guo, Y., Miller, N. R., & Bernstein, S. L. (2009). Optic nerve infarction and post-ischemic inflammation in the rodent model of anterior ischemic optic neuropathy (rAION). Brain Research, 1264, 67–75. 10.1016/j.brainres.2008.12.075

71. Zhou, T., Zheng, Y., Sun, L., Badea, S. R., Jin, Y., Liu, Y., Rolfe, A. J., Sun, H., Wang, X., Cheng, Z., Huang, Z., Zhao, N., Sun, X., Li, J., Fan, J., Lee, C., Megraw, T. L., Wu, W., Wang, G., & Ren, Y. (2019). Microvascular endothelial cells engulf myelin debris and promote macrophage recruitment and fibrosis after neural injury. Nature Neuroscience, 22(3), 421–435. 10.1038/s41593-018-0324-9

